# T-cell receptor structures and predictive models reveal comparable alpha and beta chain structural diversity despite differing genetic complexity

**DOI:** 10.1101/2024.05.20.594940

**Authors:** Nele P. Quast, Brennan Abanades, Bora Guloglu, Vijaykumar Karuppiah, Stephen Harper, Matthew I. J. Raybould, Charlotte M. Deane

## Abstract

T-cell receptor (TCR) structures are currently under-utilised in early-stage drug discovery and repertoire-scale informatics. Here, we leverage a large dataset of solved TCR structures from Immunocore to evaluate the current state-of-the-art for TCR structure prediction, and identify which regions of the TCR remain challenging to model. Through clustering analyses and the training of a TCR-specific model capable of large-scale structure prediction, we find that the alpha chain VJ-recombined loop (CDRA3) is as structurally diverse and correspondingly difficult to predict as the beta chain VDJ-recombined loop (CDRB3). This differentiates TCR variable domain loops from the genetically analogous antibody loops and supports the conjecture that both TCR alpha and beta chains are deterministic of antigen specificity. We hypothesise that the larger number of alpha chain joining genes compared to beta chain joining genes compensates for the lack of a diversity gene segment.

Overall, our study demonstrates that valuable structure-function relationships can lie in alpha chains despite their simpler junctions. We also provide over 1.5M predicted TCR structures to enable repertoire structural analysis and elucidate strategies towards improving the accuracy of future TCR structure predictors.

## 1 Introduction

T-cell receptors (TCRs) are highly sequence diverse, somatically-recombined receptor proteins. They govern the T-cell mediated adaptive immune response by interacting with antigens such as peptides presented by the Major Histocompatibility Complex (pMHC) [1–3]. The potential of TCRs as therapeutics for cancer and other diseases has led to substantial research effort dedicated to understanding and predicting their pMHC interactions and specificity e.g. [4–19]. Despite the importance of this task [20], TCR complementarity remains extremely difficult to predict beyond well-explored case studies, with existing computational models failing to generalise to unseen data [21, 22]. The development of more generalisable approaches was recently labelled a ‘Cancer Grand Challenge’ by the US National Institutes of Health [23].

Currently, most TCR:pMHC specificity predictors use sequence based features only [4–18]. The incorporation of structural information offers a compelling strategy to encourage models to learn more generalisable physicochemical principles of complementarity [24–26]. However, accessing and generating structural information for TCR:pMHC specificity prediction poses new challenges, which is perhaps why only a handful of structure-aware methods exist to date [24, 27]. One bottleneck is the low quantity of available TCR structure data and the high experimental cost of obtaining more: only 605 publicly available crystal structures deposited in the PDB [28] at the time this study was conducted contained TCR structures [29], compared to the over 20,000 epitope-labelled paired TCR sequences available in VDJdb [30]. However, advances in computational TCR structure prediction provide a cheaper and higher throughput alternative to experimentally solving TCR structures. While historically the accuracy of adaptive immune receptor structure prediction algorithms has been poor [31–33], recent progress applying deep learning methods has made faster and higher-accuracy *in-silico* predictions of TCR structures viable [34, 35]. Simultaneously, federated learning initiatives are increasing accessibility to highly valuable datasets from the Pharmaceutical industry, which could potentially significantly boost the amount of available data for model training [36, 37]. A second limitation of many TCR:pMHC specificity predictors is their inclusion of only beta chain information of the TCR [4, 7, 8, 11, 12, 21, 38, 39]. TCRs function as heterodimers, typically consisting of an alpha and a beta chain whose recombination and sequence diversification process relies on the same genetic mechanisms underpinning the synthesis of the heavy and light chain of näıve B-cell receptors (BCRs). Briefly, sequence diversity results from the recombination of multiple genes (VDJ for beta/heavy chains, VJ for alpha/light chains) leading to six structurally proximal complementarity determining region (CDR), loops, three per chain [3]. While the CDR1 and CDR2 loops are relatively less sequence variable because they are templated by the V-genes, the CDR3 loops lie across the gene junctions giving rise to combinatorial diversity in their sequence. Further Nand Pnucleotide insertions and deletions at the junctions in the CDR3 lead to substantially greater length and sequence diversity [40]. Overall, it is understood that the VDJ-recombined beta/heavy chain CDR3 (CDRB3/CDRH3) loop exhibits the greatest sequence diversity and thus contributes more to determining target specificity than the corresponding VJ-recombined CDR3 of the alpha/light chain (CDRA3/CDRL3) [3, 41, 42]. However, while there is strong evidence that this holds for BCRs/antibodies, where the CDRH3 loop appears to contribute disproportionately to antigen specificity relative to the CDRL3 loop [43, 44], it remains unclear whether the TCR CDRB3 is more deterministic of antigen recognition than the CDRA3 [45], implying that the exclusion of the alpha chain could lead to the loss of important information for specificity prediction.

Here, leveraging a proprietary dataset of TCR crystal structures solved by Immunocore to supplement the publicly available data, we evaluated the current headline performance of computational TCR modelling and identified which CDRs remain challenging to predict, representing barriers to the utility of structural models for predicting properties of TCRs such as pMHC specificity. We first clustered the complete structural dataset to identify relevant biases, which unexpectedly revealed that the CDRA3 and CDRB3 loops exhibit similar structural diversity, despite their differing recombination mechanisms.

We then built TCRBuilder2+ by retraining TCRBuilder2 [35], a best-in-class TCR specific structure predictor, on the expanded dataset of structures. The performance of TCRBuilder2+ improved for genes better sampled in the new training set, and TCRBuilder2+ achieves comparable overall accuracy to Alphafold-Multimer at a fraction of the computational cost. Consistent with the clustering analysis, both TCRBuilder2+ and Alphafold Multimer found the VJ-recombined CDRA3 loop at least as challenging to model as the corresponding VDJ-recombined CDRB3 loop, differentiating TCR structure prediction from BCR/antibody structure prediction, where the CDRL3 is substantially easier to predict than the CDRH3 [35, 46]. In order to further explore the structural diversity of TCRs we used TCRBuilder2+ to predict the structures of over 1.5 million TCR sequences from the Observed T-cell receptor Space (OTS) database [47].

Finally, we investigated potential sources of the high structural diversity in the CDRA3. The amino acid composition and genetic coherence of distinct structures suggest that the greater number of TRAJ genes relative to both TRBJ and IGKJ/LJ genes, which leads to greater combinatorial diversity in the alpha chain, may also lead to enhanced structural variability.

Our study is the first to elucidate the surprisingly high structural complexity of the TCR CDRA3 and the challenge of predicting its structure, and reinforces the value of paired-chain information for capturing the function of TCRs.

## 2 Results

### 2.1 Curating and analysing an industry-augmented dataset of solved TCR structures

The amount of experimentally-determined TCR structure data in the public domain is significantly less than TCR sequence data [29], therefore harnessing structural features in general TCR property prediction pipelines requires the ability to make accurate, and ideally fast, predictions of TCR structures. In recent years, deep learning methods have emerged as state of the art for this task [35]. However, all machine learning models are limited by the data available to train them on and there are countless examples of deep learning models that generalise poorly or that learn dataset biases rather than the underlying function governing the phenomenon [22, 48–50].

In order to explore how increased structural data could improve TCR structure prediction, we supplemented the publicly available data with an internal repository of TCR structures solved by Immunocore, retraining our latest high-throughput TCR-specific deep learning model, TCRBuilder2 [35] (Fig. 1a). As an initial step we analysed the novel dataset of TCR structures to identify any residual or newly introduced data biases and to enable fairer downstream model evaluation.

**Figure 1:**
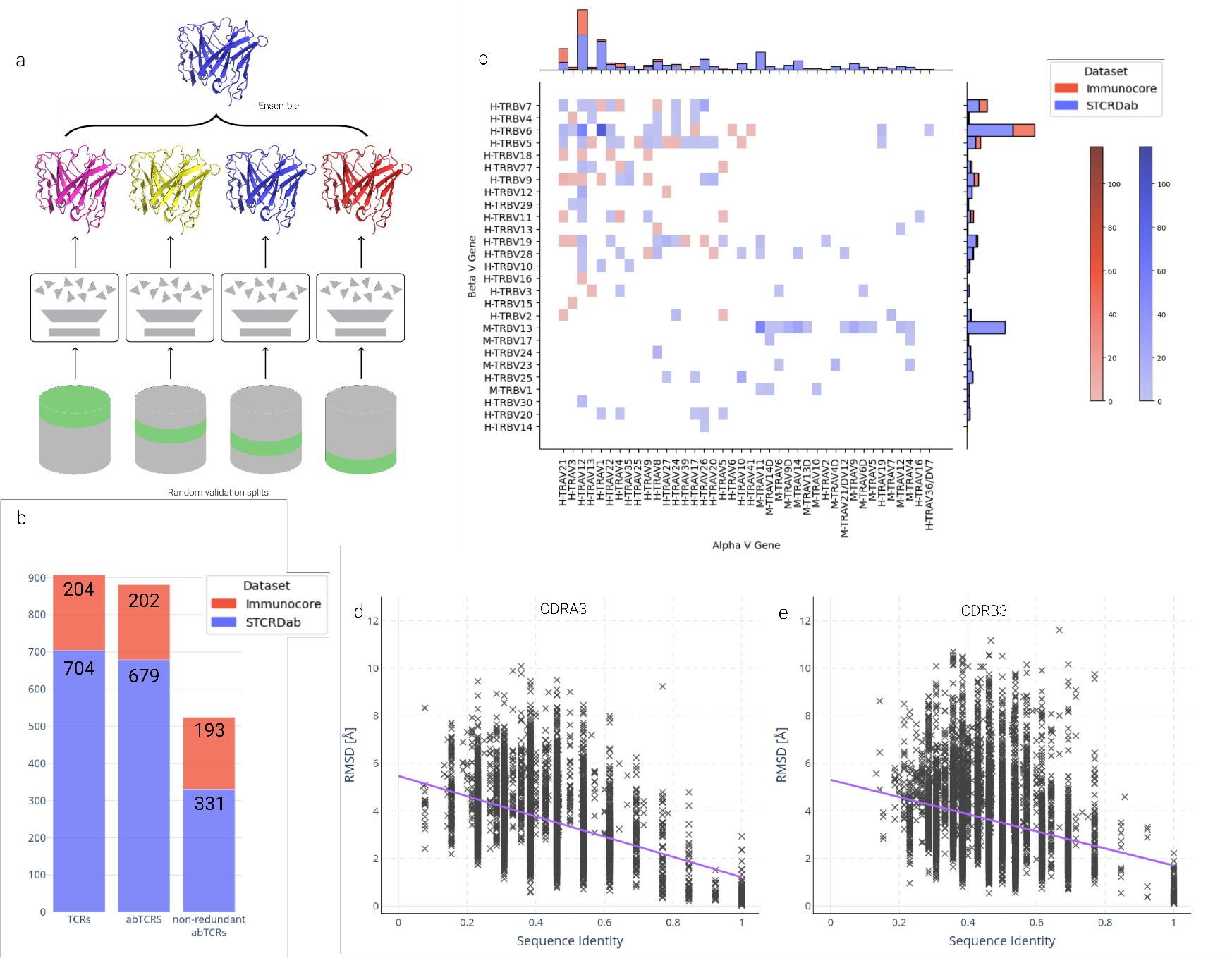
Overview of the architecture and training data distribution of TCRBuilder2+. a) TCRBuilder2 architecture [35] for TCR structure prediction. An ensemble of models is trained using different training / validation splits to generate four predictions per sequence. The final model is chosen as the model closest to the mean prediction of the ensemble. b) The joint distribution over paired alpha and beta V genes of TCR structures in the training set. The paired space is sparsely sampled. c) Adding Immunocore data to the STCRDab data expands the training set by 204 structures, and increases the number of non-redundant sequences by 58%. Genes beginning with ‘H-’ are human, those beginning with ‘M-’ are murine. d&e) Pairwise RMSD of CDRA3 (d) and CDRB3 (e) loops in the training data against pairwise sequence identity with a linear trend fit (purple, (*PCC_CDRA_*_3_ = *−*0.41, *PCC_CDRB_*_3_ = *−*0.27)). For sequence identities *>* 0.65, sequence identity and RMSD of the CDR3 loops is somewhat correlated (*PCC_CDRA_*_3_ = *−*0.66, *PCC_CDRB_*_3_ = *−*0.52). Sequence identities *<* 0.65 are substantially less correlated with RMSD for both CDR3 loops (*PCC_CDRA_*_3_ = *−*0.29, *PCC_CDRB_*_3_ = *−*0.17). CDRB3 displays a slightly greater range of RMSD than CDRA3 loops.

The original training data for TCRBuilder2 [35] contained 704 experimentally resolved TCR structures sourced from STCRDab [29, 51]. These structures were obtained from 463 distinct PDB files, with 349 of the 704 structures having a unique sequence. Our novel data contained 204 Immunocore TCR structures, of which 201 sequences are unique and only 6 overlap with the existing training set (Fig. 1b). This expansion contributed to a combined set of 908 structures, of which 544 are unique in sequence. Filtering for alpha-beta TCR pairs resulted in 881 TCR structures to train TCRBuilder2+, of which 524 are sequence non-redundant. Furthermore, since the original TCRBuilder2 was published more TCR structures have become publicly available, allowing for the curation of a larger and more robust test set. We generated a new, independent, and non-redundant test set of 45 alpha-beta TCR structures for downstream benchmarking (SI Tab. 1).

**Table 1:** Mean RMSD of TCR structure predictions by different prediction models evaluated on the test set. *LYRA, the homology method, could only be evaluated on 38 of the 45 test structures for which a structure was generated. The best mean RMSD for any TCR region is emboldened, the second best is italicised. Backbone RMSD is reported for each region of the TCR. A/B the entire alpha and beta chain respectively, FWA/FWB the constant framework region of each chain, CDRA/CDRB the three CDR loops of each chain.

|  | AlphaFold Multimer [56] | TCRBuilder2+ | TCRBuilder2 [35] | LYRA* [33] | ABodyBuilder2 [35] |
| --- | --- | --- | --- | --- | --- |
| A | **1.02** | 1.20 | *1.16* | 1.19 | 3.51 |
| FWA | **0.72** | 0.99 | 0.92 | *0.79* | 3.00 |
| CDRA1 | **1.11** | *1.25* | 1.26 | 1.51 | 3.62 |
| CDRA2 | 0.93 | 0.96 | *0.83* | **0.81** | 7.03 |
| CDRA3 | **2.00** | *2.06* | *2.06* | 2.69 | 3.29 |
| B | **0.95** | **0.95** | 0.96 | 0.99 | 3.30 |
| FWB | 0.74 | 0.73 | **0.71** | **0.71** | 3.18 |
| CDRB1 | *0.66* | 0.75 | **0.64** | 0.78 | 3.42 |
| CDRB2 | **0.59** | 0.78 | *0.69* | 0.74 | 3.65 |
| CDRB3 | **1.70** | *1.78* | 1.92 | 2.27 | 3.42 |

Figure 1c shows the distribution of TRAV and TRBV genes of the 881 TCR structures contained in our new training set, characterising the biases present in the data. There are strong biases towards a small number of genes and overall the joint distribution over pairs is sparsely sampled, with the majority of alpha/beta pairs unsampled or only appearing in the dataset once. The joint distribution is disproportionately weighted towards a small subset of gene pairs; for example, TRAV12 and TRAV1 paired with TRBV6 account for 212 structures, 24.1% of the training data. Observations of naturally occurring alpha/beta chain pairings imply variable gene pairings are relatively uniform after accounting for gene abundance [47], indicating that it is the selection of TCR structures that have been experimentally resolved that is giving rise to the TRAV and TRBV gene bias, not the natural underlying TCR distribution. To disentangle the contributions of the STCRDab and the Immunocore datasets to the bias we also considered the distributions independently (SI Fig. 1). While we observed similar biases within each set, our new dataset improves the sampling of human TRAV-TRBV pairs. We saw a similar trend of bias and improved coverage in our new dataset for TRAJ-TRBJ pairs, as well as VJ pairings, shown in SI Figures 2-4.

**Figure 2:**
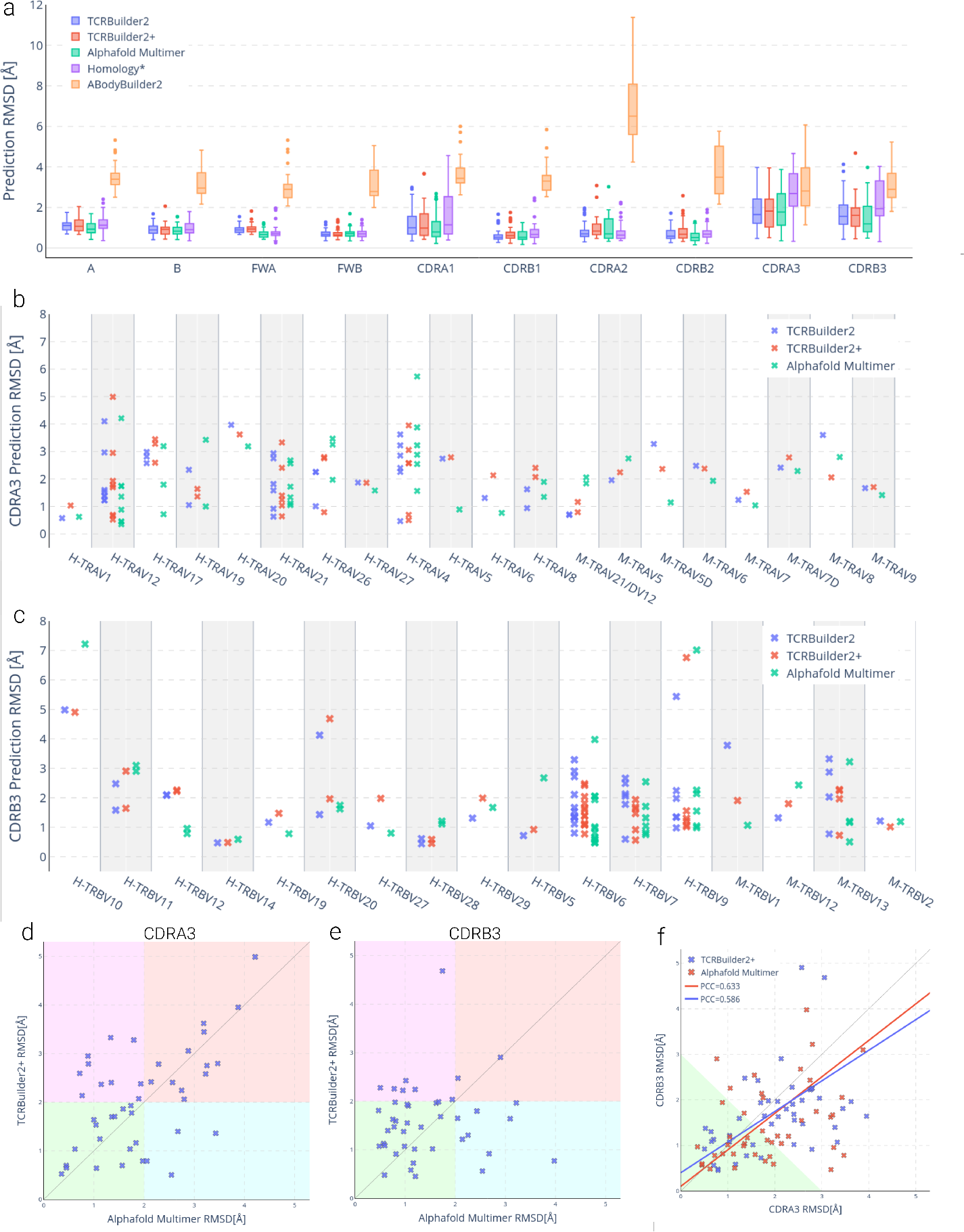
Evaluation of TCR structure prediction models on our test set. (a) Structure prediction RMSD by TCR region. (b&c) Structure prediction accuracy over (b) CDRA3 by TRAV gene, and (c) CDRB3 by TRBV gene for TCRBuilder2 (blue), TCRBuilder2+ (red) and Alphafold Multimer (green). Genes beginning with ‘H-’ are human, those beginning with ‘M-’ are murine. (d&e) TCRBuilder2+ vs Alphafold Multimer prediction RMSD of CDRA3 (d) and CDRB3 (e). Points in the green quadrant or red quadrant indicate both methods are able to (green) or unable to (red) predict the structure to within 2Å, respectively, while points in the purple quadrant or cyan quadrant indicate that Alphafold-Multimer (purple) or TCRBuilder2+ (cyan) is uniquely able to model the loop to within 2Å. (f) CDRB3 vs CDRA3 prediction RMSD. For reference, subsequent docking case studies suggest that predictions within the green shaded region (CDRA3 RMSD + CDRB3 RMSD *<* 3Å) can be reliably docked with an interface RMSD *<* 4Å (see Section 2.4, Fig. 12a).

Computationally predicting TCR structures from their sequences using data-driven methods such as deep learning or homology relies on there being a functional mapping from sequence to structure that can be extracted from the existing data. Therefore, we used the supplemented structural data to investigate the relationship between sequence and structure identity of most variable TCR loops, the CDRA3 and CDRB3 (Figs. 1d-1e).

There is a disparity in trend between sequence identities of CDR3 loops greater than 65% and between 0% and 65%. At lower sequence identities, structural distances are weakly correlated (PCC -0.17 to -0.29). Higher sequence identities are moderately correlated with smaller structural distances (PCC -0.52 to -0.66), though, even between loops of identical sequence, substantial structural deviation can be observed: CDRB3 and CDRA3 loops with very high sequence identities (*≥* 90%) can display structural dissimilarities of up to 4Å. These results demonstrate the known limitations of homology approaches to TCR structure prediction, as there is often more complexity in what determines CDR3 structure than sequence identity of the loop alone. Deep learning approaches, such as the TCRBuilder2 architecture, have been shown empirically to be able to capture more subtle and complex patterns from data, yielding improvements in predictive capability [34, 35, 46]. The sequence identity to structural distance distributions in Figures 1d and 1e also provide useful background distributions against which to evaluate the predictive performance of TCR structure prediction methods, since they provide an upper bound of the expected RMSD for a given prediction if a template with the closest sequence identity to the query sequence were selected. We observed a wide range of structural distances between CDR3 loops across all sequence identities, many of which are substantially greater than 2.0Å.

Finally, comparing the pairwise RMSD distributions of the CDRA3 and CDRB3 loops reveals that the CDRA3 loop (Fig. 1d) is as inherently structurally diverse as the CDRB3 loop (Fig. 1e, SI Fig. 6). We confirmed this finding by clustering each of the CDR3 loop types by RMSD (SI Tab. 2). Surprisingly, despite their simpler genetic composition CDRA3 loops do not cluster into fewer structural clusters than CDRB3 loops; by contrast, the CDRL3 loops of antibodies cluster into far fewer shapes than their CDRH3 counterparts [52–55]. This disparity between TCR and antibody structures hints that the structures adopted by CDRA3 loops may be more complex than suggested by their genetic recombination mechanism alone.

**Table 2:** Definitions for anchors and loops based on IMGT numbering. Structures were aligned by anchors and RMSD was then calculated over residues in the loops.

| Region | Anchor residue numbers | Loop residue numbers |
| --- | --- | --- |
| CDR1 | 22-27, 39-44 | 28-38 |
| CDR2 | 51-56, 66-71 | 57-65 |
| CDR3 | 100-105, 118-123 | 106-117 |

### 2.2 Benchmarking state-of-the-art TCR structure prediction models

We retrained TCRBuilder2 on our supplemented dataset to generate a new model (‘TCRBuilder2+’), capturing the current performance ceiling of a TCR-specific deep learning model for high-throughput structure prediction. While the original TCRBuilder2 model followed from ABodyBuilder2 [35] and selected the ensemble of trained models based on only the beta chain validation set accuracy, in this work we adapted the ensemble selection process to include the accuracy over both chains. This adaptation better reflects TCR biology by accounting for the high structural diversity we observed in the alpha chain.

We benchmarked TCRBuilder2+ against the original TCRBuilder2 [35], a TCR-specific homology modeller (LYRA [33]), an antibody-specific deep learning model (ABodyBuilder2 [35]), and the state-of-the-art general protein structure predictor (Alphafold-Multimer [56]). We report the mean RMSD and the RMSD distribution of the 45 test set structures predicted by different models in Table 1 and Figure 2a, respectively.

As expected, predicting TCR structures with ABodyBuilder2 yields low accuracy predictions across all TCR regions despite the global similarities between TCRs and antibodies, confirming that structure prediction models trained on protein family subsets learn to make accurate predictions within the distribution of the data they are trained with, rather than generalising beyond it [57]. The homology model LYRA [33] only produces predictions for sequences for which a suitable template is found, and so we are only able to benchmark 38 of the 45 test structures. We provide a comparison for those 38 predictions only in SI Figure 7. Overall, homology modelling performs competitively for the framework regions (FW) and the germline encoded CDRA2, CDRB1 and CDRB2, but struggles with the CDRA3, CDRB3, and CDRA1 loops.

TCRBuilder2, TCRBuilder2+ and Alphafold-Multimer all exhibit competitive performance across the whole test set. TCRBuilder2+ outperforms TCRBuilder2 on CDRB3, but performs similarly across the other regions (Tab. 1), suggesting that the supplemented training dataset only yields limited average improvements. Based on the median RMSD, TCRBuilder2+ prediction accuracy is on par with TCRBuilder2 and is only slightly outperformed by Alphafold-Multimer (Fig. 2a). Examining how frequently each method makes a prediction of reasonable accuracy (*<* 2Å, see Methods), we found that the framework regions (FW) and germline-encoded loops (CDR1 and CDR2) are largely well predicted. As was observed for the homology model, CDRA1 is markedly more difficult to predict than CDRB1, with mean TCRBuilder2+ prediction RMSDs of 1.25*±*0.81Å and 0.75 *±* 0.41Å respectively. Meanwhile, the CDRB3 and CDRA3 loop predictions exhibit the largest errors, as is expected due to their greater sequence and structure diversity relative to the germline-encoded loops. Specifically, TCRBuilder2+ predicts 34 (75.6%) CDRB3 structures with sub-2Å RMSD, more than TCRBuilder2 and Alphafold-Multimer which predict 26 (57.8%) and 32 (71.1%) within that threshold, respectively (SI Tab. 3). On the other hand, Alphafold-Multimer is able to predict 27 (60.0%) of CDRA3 structures within 2Å RMSD, whereas TCRBuilder2+ and TCRBuilder2 respectively retrieve 21 (46.6%) and 23 (51.1%) CDRA3 structures within that threshold (SI Tab. 3). Both the general method (Alphafold-Multimer) and TCRBuilder2+ predict CDRB3 loops with a higher average accuracy than CDRA3 loops (Tab. 1, Fig. 2a), once again contrary to the conjecture that the greater junctional diversity of CDRB3 compared to CDRA3 ought to lead to higher structural complexity.

Due to the biases towards specific gene pairings in the training set (Fig. 1c), we then analysed the CDRA3 and CDRB3 predictive accuracy over different V-genes in the test set (Figs. 2b-2c). Constrasting the two TCR-specific models, we find that predictive accuracy improves for genes that are better represented in the new training set (Fig. 1c, SI Fig. 8-9), with TCRBuilder2+ resulting in, on average, higher accuracy CDRA3 predictions than TCRBuilder2 for human TRAV19 and -21 and CDRB3 predictions for human TRBV6, -7, and -9. This suggests that more data for a specific gene helps predict the structures of TCRs that derive from that gene, but that this improved performance might not necessarily translate to all TCRs generally. This in turn implies that increasing the amount and improving the genetic sampling of experimentally solved structure data with which models can be trained will be necessary to improve general inference of TCR structures.

To assess whether TCR-specific and general structure predictors struggle to model the same TCR sequences, we compared the RMSD of the TCRBuilder2+ and Alphafold-Multimer CDRA3 and CDRB3 loop predictions for every test structure (Figs. 2d-2e). Points in the green quadrant or red quadrant indicate both methods are able to (green) or unable to (red) predict the structure to within 2Å, while points in the purple quadrant or cyan quadrant indicate that Alphafold-Multimer (purple) or TCRBuilder2+ (cyan) is uniquely able to model the loop to within 2Å. We observe for CDRA3 loops that a substantial number (13/45) fall into the red quadrant, indicating that neither TCRBuilder2+ nor Alphafold-Multimer can accurately predict their conformation. In contrast, very few CDRB3 loops fall into the red quadrant (2/45), reiterating that CDRA3 structure is more challenging to predict regardless of the model used.

A hypothetical ensemble model able to perfectly identify whether the TCRBuilder2+ or AlphafoldMultimer prediction is a better CDRB3 or CDRA3 loop model — independent of the other loop — would currently achieve 95.6% under 2Å for CDRB3 (up from 75.6% for TCRBuilder2+ alone) or 71.1% under 2Å for CDRA3 (up from 60.0% for Alphafold-Multimer alone). This suggests that a method that can selectively incorporate knowledge of general protein loop conformations into TCR loop structure prediction may translate to larger performance boosts for CDRB3 than CDRA3. Furthermore, the fact that even when combining the best predictions of both TCRBuilder2+ and Alphafold-Multimer the CDRA3 structure is more elusive to predict emphasises its structural complexity despite its simpler genetic recombination.

As an indication of whether deep learning approaches have learned to model the TCR variable region globally rather than considering loops independently of one another, we investigated the correlation between CDRA3 and CDRB3 predictive accuracy (Fig. 2f). The Pearson’s correlation coefficients (PCC) calculated for both Alphafold Multimer (0.633) and TCRBuilder2+ (0.586) indicate a slight correlation between CDRA3 and CDRB3 prediction RMSD, suggesting that deep learning models consider loop conformations somewhat cooperatively. While comparing the RMSD of the CDRA3 and CDRB3 of each TCR in the test set, we found that 64.4% (29/45) and 60% (27/45) of TCRBuilder2+ and Alphafold Multimer’s predictions respectively fall below the RMSD identity line (Fig. 2f), confirming that for the majority of test structures the CDRA3 is more difficult to model than the CDRB3 of a given TCR, regardless which model is used.

Finally, we explored whether TCR structural models have yet attained a level of accuracy where useful information can be extracted from TCR:pMHC interfaces predicted using physics-based docking [58] and present the extended results of these experiments in the Supplementary Information (SI Sec. 3). We found that a total RMSD over both the CDRA3 and CDRB3 *<* 3Å strongly supports the retrieval of an accurate docked pose (interface RMSD over all heavy atoms *<* 4Å in all cases we ran simulations for, indicating that structure predictions are beginning to show merit for TCR:pMHC interface prediction. Our evaluation of current state of the art structure prediction models suggests a third of TCRs are modelled within this threshold (green region in Figure 2f), indicating further improvements are required to enable consistent interface prediction *via* docking. It is not possible to identify a consistent accuracy threshold for docking when considering the CDRA3 or CDRB3 individually (SI Fig. 11), highlighting once again the importance of accurately modeling both regions for TCR:pMHC interface prediction.

In summary, retraining TCRBuilder2 with substantially more training data did not yield large gains in average predictive accuracy, although the mean RMSD for CDRB3 reduced from 1.92Å (TCRBuilder2) to 1.78Å (TCRBuilder2+), narrowing the gap to Alphafold-Multimer’s benchmark of 1.70Å. We did observe improvements across TRAV and TRBV genes for which the abundance of training examples increased, suggesting that increasing the diversity of training examples by experimentally solving as yet unsampled genes should yield accuracy improvements. Physics-based docking studies indicate that, while predicted TCR structures are beginning to show utility for predicting TCR:pMHC interfaces, improvements in TCR structure prediction accuracy should also yield improvements in docking. Furthermore, and in-line with our observations of experimentally solved TCR structures, we found that the CDRA3 is consistently more challenging to predict than the CDRB3, with evaluations of all models suggesting that the genetically simpler CDR3 junction region, CDRA3, remains a key barrier to accurate TCR modelling.

### 2.3 Enabling structure-based deep learning research with 1.5 million TCR structure predictions

Next, we estimated to what degree the apparent diversity of alpha chain CDR structures is sampled by natural TCR repertoires. To assess this, we harnessed the computational efficiency of TCRBuilder2+ and predicted the structures of over 1.56M non-redundant TCR sequences from the newly released OTS database [47], yielding an approximately 2500-fold increase in TCR structures in the public domain. We then conducted a structural analysis, contrasting the TCRs in OTS to the BCRs/antibodies from OAS [59]. Our previous analysis of crystal structures suggested that the CDRA3 of TCRs is as structurally diverse as the CDRB3; here, we observe that TCR CDRA3 loops from T cell repertoire sequencing also exhibit this diversity.

Figures 3a & b show that the CDR1 and CDR2 loops of (a) the TCR alpha and beta chains, and (b) the antibody light and heavy chains cluster into canonical forms, as has been shown to be the case for experimentally solved structures [60, 61]. Specifically, we find that for 1000 randomly sampled TCRs from our OTS structure predictions, the CDRA1, CDRA2, CDRB1, and CDRB2 loops fall into 126, 200, 34, and 105 clusters, respectively, when applying a 1Å RMSD threshold (SI Tab. 5). Repeating this clustering with 1000 antibodies from OAS, we find that the CDRL1, CDRL2, CDRH1, and CDRH2 separate into 31, 9, 19, and 36 distinct loop conformations (SI Tab. 6).

**Figure 3:**
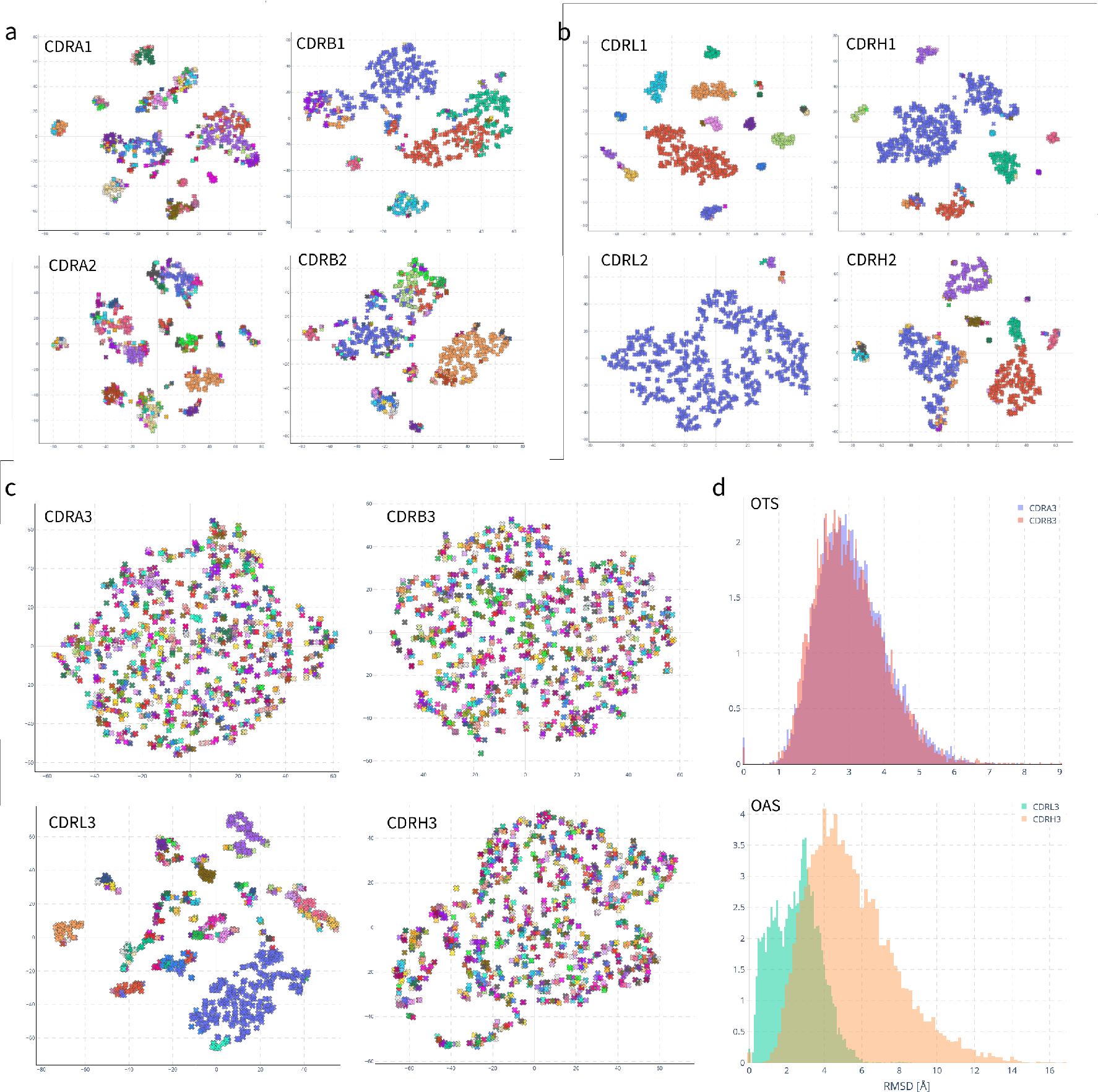
Structure distributions of 1000 randomly sampled TCRBuilder2+ predictions from the OTS database [47] and 1000 randomly sampled ABodyBuilder2 [35] predictions from the OAS database [59]. All t-SNE plots are coloured by clustering with a 1Å RMSD threshold. a & b) t-SNE clustering of RMSD between canonical loops of TCRs (a) and antibodies (b). CDR1and 2 visibly cluster into canonical forms. c) t-SNE clustering of RMSD between TCR CDRA3 and CDRB3, and antibody CDRL3 and CDRH3. The antibody CDRL3 loop clusters into canonical forms, whereas the TCR CDRA3 does not. No clustering is observed for the TCR CDRA3, CDRB3 or the antibody CDRH3. d) Distribution of pairwise RMSD between CDRA3 and CDRB3 predictions of OTS and between CDRL3 and CDRH3 predictions of OAS. The distribution of pairwise RMSD of predicted TCRs is indistinguishable between CDRA3 and CDRB3, demonstrating equivalent structural diversity in natural repertoires. In antibodies the pairwise RMSD distribution of CDRL3 is shifted towards lower RMSD than that of CDRH3, indicative of CDRL3 structures with high structural similarity, as would be expected of canonical loops.

**Figure 4:**
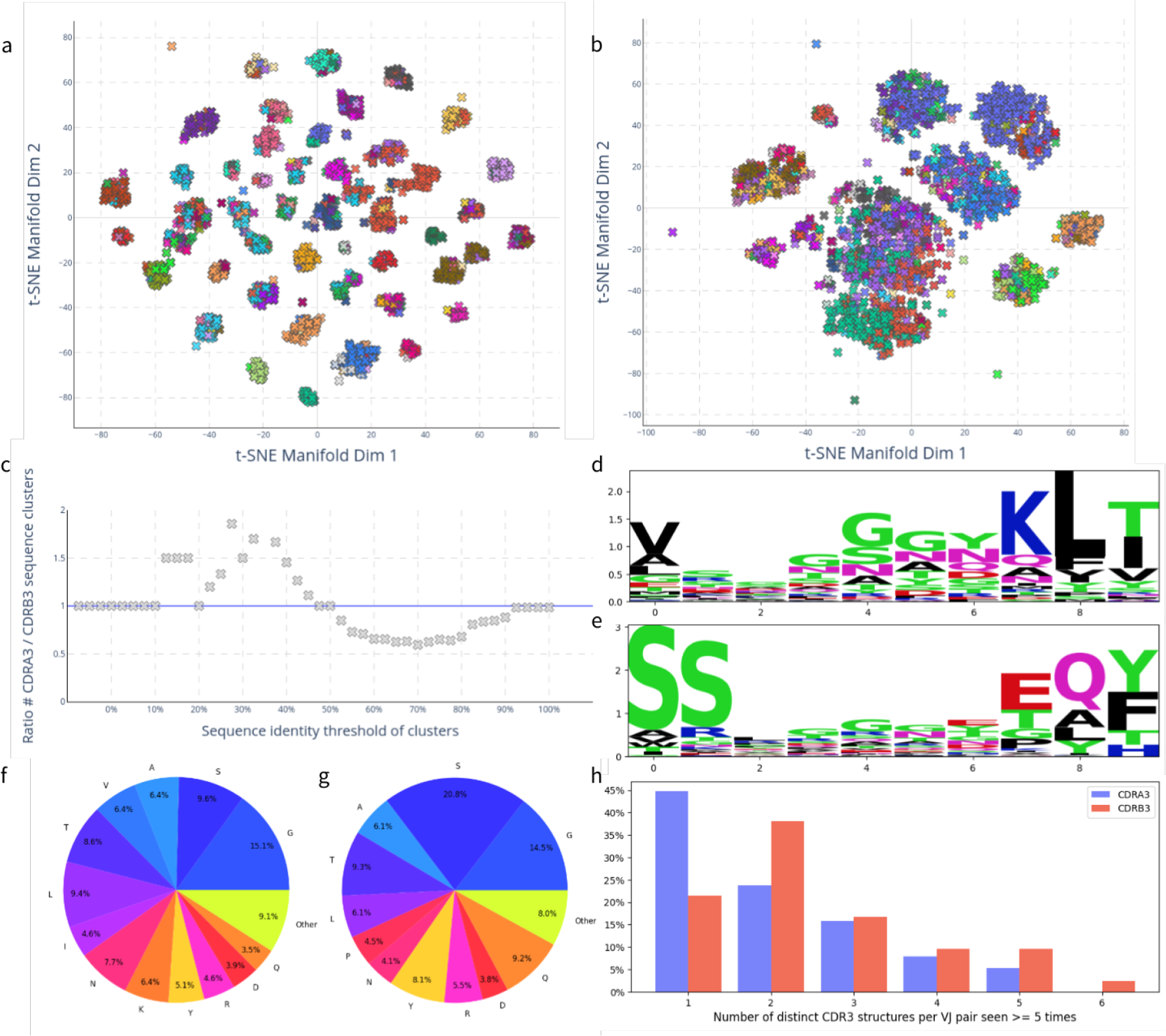
a & b) Sequence identity t-SNE distributions of 2000 OTS (a) CDRA3 and (b) CDRB3 loops coloured by 50% sequence identity cluster. CDRA3 sequences separate into fewer and more distinct clusters. c) Ratio of CDRA3 sequence clusters to CDRB3 sequence clusters varied by sequence identity threshold. At high sequence identities more CDRB3 clusters emerge, whereas at lower sequence identities more CDRA3 clusters emerge. d & e) Sequence logos of 2000 OTS samples’ (d) CDRA3 and (e) CDRB3 loops of length ten. Central regions of the CDRA3 and CDRB3 sequence have high entropy, and reveal high proportions of glycine. f & g) Distribution of amino acids across 2000 OTS (f) CDRA3 and (g) CDRB3 samples. Blue-tinted segments indicate relatively small amino acids which may enable loop flexibility. h) Histogram of the number of 2Å structural clusters for variable (V) and joining (J) gene pairs (TRAV & TRAJ for CDRA3 and TRBV & TRBJ for CDRB3) in training data structures for which at least five structures exist per VJ pair. VJ gene pairings map onto unique CDRA3 structures in 44% of cases and onto unique CDRB3 structures in 22% of cases.

In agreement with our analysis of experimentally solved TCR structures, neither the CDRA3 nor CDRB3 of naturally occurring OTS TCRs cluster into distinct structures: 1000 random TCRs fall into 840 and 828 clusters for CDRA3 and CDRB3 respectively (Fig. 3c). As expected, the antibody CDRH3 also does not cluster into distinct shapes (880 clusters per 1000 antibodies), but the CDRL3 loop does (131 clusters per 1000 antibodies). Comparing the clustering of antibody CDRL3 to TCR CDRA3 structures exemplifies the increased structural diversity of the CDRA3 despite both loop sequences being the product of only VJ recombination.

The pairwise RMSD distributions of the CDRA3 and CDRB3 of the OTS predictions are indistinguishable (Fig. 3d), confirming that TCRBuilder2+ generates equally diverse conformations of these regions when predicting sequences in OTS. On the other hand, the RMSD distribution of the antibody CDRL3 is shifted towards lower RMSDs relative to CDRH3 and exhibits the bimodality expected of loops sampled from distinct canonical shapes; this bimodality is not observed for the TCR CDRA3.

Together, this analysis of predicted TCR and antibody structures demonstrates the difference in structural diversity of the VJ recombined CDRA3 and CDRL3 loops. This concurs with our finding from the analysis of TCR crystal structures and the evaluation of TCR prediction models that CDRA3 structural complexity matches that of CDRB3.

### 2.4 Potential explanations for CDRA3 structure diversity

Finally, we sought an explanation for the unexpectedly high structural diversity of CDRA3 by investigating the CDRA3 and CDRB3 sequence distributions (Fig. 4). Unlike the CDRB3 sequences, CDRA3 sequences cluster by sequence identity (Figs. 4a-4b). Greedy clustering confirms this: fewer CDRA3 than CDRB3 clusters emerge for sequence thresholds between 50% and 100% (Fig. 4c, SI. Tab. 9), suggesting that CDRA3 sequences can be separated into sets of similar sequences. To expand on this observation, we investigated the sequence logos of the CDRA3 (Fig. 4d) and CDRB3 (Fig. 4e), finding as expected that the central regions of both loops are highly diverse. However, the variability of the central CDRB3 residues is greater than that of the central CDRA3 residues, implying that diversity emerging from VDJ recombination with the additional junctions relative to the VJ recombination does increase the sequence diversity of the CDRB3 relative to CDRA3. Both the clustering and the slightly lower variability of the CDRA3 sequences contrasts with our observations of CDRA3 and CDRB3 structures, where CDRA3 exhibits similar diversity to CDRB3.

Both the CDRA3 and CDRB3 loop sequences have high glycine and serine content (Fig. 4d & e). Following from this observation we investigated the proportion of residues that could be conducive to flexibility or the emergence of multiple conformations per sequence, which could provide a potential explanation for structural diversity. Figures 4f & 4g suggest that the proportion of small residues conducive to loop flexibility is approximately 50% for both the CDRA3 (f) and CDRB3 (g). Furthermore, studies of TCR structure [62–64] and *in-silico* simulations [64, 65] have suggested that both the CDRA3 and CDRB3 structures may be flexible. Though it is difficult to analyse robustly computationally, we explored this hypothesis *via* a preliminary molecular dynamics case study of an apo TCR, the results of which we provide in the Supplementary Information (SI Sec. 4). In our case study we found that an array of conformations of the CDRA3 and CDRB3 loops predicted by TCRBuilder2+ and Alphafold-Multimer appear to be physically plausible local energy minima of the simulations (SI Fig. 15). Therefore, while both the amino acid composition and the molecular dynamics simulation support the possibility that both CDRA3 and CDRB3 loops are flexible, it is not possible to distinguish CDRA3 from CDRB3 or suppose that one is more or less rigid than the other.

To further explore whether the emergence of multiple conformations drives similar structural diversity despite reduced sequence diversity of the CDRA3 relative to the CDRB3, we investigated how deterministic TRA or TRB VJ gene pairing is of CDRA3 or CDRB3 loop structure, respectively (Fig. 4h). For all TRAV-TRAJ and TRBV-TRBJ combinations for which at least five structures were observed in the TCRBuilder2+ training data, we calculated how many distinct CDRA3 and CDRB3 structures existed, respectively, using a 2Å RMSD threshold. We observed that 44% of TRAV-TRAJ pairs map onto a single CDRA3 structure, while only 22% of TRBV-TRBJ pairs map onto a single CDRB3 structure. This suggests, in agreement with the expected effect of the insertion of the diversity gene and the additional junction in the CDRB3 loop, that the VJ gene pair is less deterministic of CDRB3 than CDRA3 structure. The high structural coherence of TRAV-TRAJ combination also suggests that CDRA3 structures are relatively well-defined for a given gene pair.

However, the reduced sequence complexity of the CDRA3 relative to the CDRB3 and high coherence of CDRA3 structure to TRAV-TRAJ gene pair raises the question where the equivalent CDRA3 and CDRB3 structural diversity we have observed originates from. Next, we investigated the combinatorial diversity of the CDRA3 and CDRB3 that arises from their genetic recombination mechanisms. We found that the number of potential TRAV-TRAJ pairings outnumbers those of TRBV-TRBJ (SI Tab. 7). In fact, even when accounting for the diversity gene, the combinatorial diversity of the beta chain is only about a third of that of the alpha chain, primarily due to the relatively low number of TRBJ genes compared to TRAJ. Furthermore, when clustering CDRA3 and CDRB3 sequences permissively (20-40% sequence identity), more sequence clusters emerge for CDRA3 than CDRB3 (Fig. 4c, SI Tab. 9). This suggests that CDRA3 sequences may occupy tighter but more distinct regions in sequence space, as their primary diversification occurs at the level of genetic recombination. On the other hand, CDRB3 sequences appear to exhibit greater variability at the level of individual amino acids, but are constrained by a smaller pool of genetic recombinations. It is therefore possible that CDRA3 structural diversity is driven by the larger combinatorial pairing space provided by the larger number of TRAJ genes (61) relative to TRBJ genes (14).

## 3 Discussion

Incorporating structural information into TCR specificity prediction methods offers a compelling strategy towards alleviating their current limitations [21, 22] by encouraging models to learn more generalisable physicochemical principles of complementarity *in lieu* of amino acid sequence motifs [24, 25]. Given the current sparsity of solved TCR structure data [29] and the expense of generating it, *in-silico* structure prediction is a helpful tool to provide estimates of the 3D properties of TCRs.

In this work, we have retrained a state-of-the art, high-throughput deep learning architecture for TCR structure prediction with the largest training set to date (TCRBuilder2+). We have made the resulting TCRBuilder2+ model parameters freely available (see Data Availability).

Evaluation of our model alongside Alphafold Multimer robustly characterises the current stateof-the-art of TCR structure prediction – both achieved median accuracies sub-2Å RMSD across all regions, although the CDRA3 and CDRB3 loops remain discernibly harder to predict than the other loops. While performance was similar across the whole test set, on average TCRBuilder2+ showed CDR3 predictive performance improvements for genes for which the number of examples had increased in our new training set (Figs. 2b-2c, SI Fig. 9). This aligns with related research exploring the generalisability of antibody structure prediction methods [57] as well as with more general work applying machine learning to biological data, which has shown that the quality, quantity, and sampling of the training data is critical for model performance [48, 66, 67].

We observed that a third of top-ranked TCR structure predictions exhibit sufficiently accurate CDRA3 and CDRB3 loops to enable docking within 4Å interface RMSD (Fig. 2f), implying that, while an appreciable proportion of TCR structure predictions could already be useful for physics based docking, there is still scope for improvements in prediction accuracy to benefit downstream tasks. Furthermore, disparities between poses with the best docking scores and the true best docks in our case studies suggest bespoke TCR:pMHC scoring functions may also be beneficial.

In agreement with prior findings, our simulation results suggest that the CDRs of TCRs may be flexible and may exhibit a degree of induced fit upon binding to the pMHC [62–65]. This would imply that next-generation structure predictors would benefit from information about a partner antigen to generate more complementary TCR structures, and that a hypothetical, perfect structure predictor would require information about the antigen to predict an exact single prediction. Moreover, current structure prediction models are predominantly trained on TCRs in complex with antigen, since these are considerably more abundant in the existing data [29]. If the antigen significantly influences the eventual bound structure of the TCR, then it is likely that current model predictions will struggle accurately predict apo TCR structures. While neither Alphafold-Multimer nor TCRBuilder2+ were trained to predict multiple conformations and evaluating flexibility of the predictions is challenging, both are comprised of ensembles, and as such multiple distinct predictions for a single TCR sequence can be generated (SI Fig. 14), at the expense of computational resource.

The inference speed of TCRBuilder2+ enabled the prediction of over 1.5 million sequence nonredundant TCR structures from repertoire studies, the first dataset of its kind^1^. Structurally characterising repertoire data has been shown to yield clinically relevant insights for BCRs [68–70] and has enabled the training of deep learning architectures on structural data at previously inaccessible scales [71, 72]. Our release of the OTS structure predictions has the potential to enable similar advances in computational TCR research.

We found that both TCR-specific and general protein structure predictors modelled the VJ-recombined CDRA3 loop with similar accuracy to the VDJ-recombined CDRB3 loop. Our analysis of the landscape of solved TCR structures also found that the VJ-recombined junction in the alpha chain appears to be as structurally diverse as the VDJ-recombined junction of the beta chain. This contrasts strongly with the trends seen across B cell receptors and antibodies: despite analogous CDR3 diversification mechanisms, CDRL3 loops predominantly fold into canonical structures, while CDRH3 structures are substantially more structurally diverse [35, 46, 60].

One potential functional rationale for antibodies and TCRs to have evolved these different levels of structural diversity in their VJ junction is that, in antibodies, the CDRL3 tends to plays a supportive role in binding, with the CDRH3 driving antigen specificity and binding [44], whereas current evidence suggests that the TCR CDRA3 and CDRB3 play a more equal role in pMHC binding. For antibodies this has been substantiated through recent observations that the heavy chain clonotype is predictive of binding specificity but not the light chain clonotype [43] and metadynamics studies suggesting that CDRH3s sample conformations in the space left unoccupied by the CDRL3 loop and not *vice versa* [73]. While elucidating the precise rules of TCR to pMHC binding remains an active topic of research, structural analyses have shown that the CDRA3 loop frequently makes direct peptide contact in TCR-pMHC interfaces [29, 74], and samples a similar length distribution to CDRB3 [74]. Thus, it would be coherent for evolutionary pressure to have increased CDRA3 structural diversity alongside CDRB3 diversity to achieve better peptide sensitivity.

Supportive of this hypothesis, the number of human TRA and TRB locus genes indicate that the potential combinatorial diversity of the alpha chain is greater than that of the beta chain, compensating somewhat for the presence of only one junction region rather than two (SI Tab. 7 [75]). This appears to be driven primarily by the greater number of TRAJ genes (61) relative to both TRBJ (14) and IGLJ/KJ (11/5). Alpha chain combinatorial and structural diversity substantially exceeds that of the antibody light chain, which remains less structurally diverse despite undergoing additional somatic hypermutation diversification. Overall, this outsized diversity in the alpha chain VJ junction, coupled with relative data paucity, likely contributes substantially towards the CDRA3 prediction errors being on-par with those of the CDRB3 across all models tested. However, the high structural coherence of TRAJ-TRAV gene pairs and observed predictive improvements over better sampled genes suggests that supplementing existing TCR structure data with as yet unsolved crystal structures could yield substantial improvements in prediction accuracy.

TCRBuilder2+ allows TCR structures to be predicted at high-throughput with sub-2Å median RMSD across all regions, but near-term improvements on these values may require fundamental methodological changes that better reflect TCR biology. Our analysis highlights the structural variability of the TCR CDRA3, on par with that of the CDRB3, differentiating T cells from their B cell/antibody cousins and reinforcing the value of paired-chain sequencing data and incorporating both chains in TCR:pMHC specificity models. Finally, our industry-augmented high-throughput model, TCRBuilder2+, enables the integration of TCR structure predictions into *in-silico* TCR screening and design pipelines, and, in combination with our release of ca. 1.5 million predicted structures of native TCRs, will help unlock the field of repertoire structural immunoinformatics.

## 4 Methods

### 4.1 Data curation

The original training data of TCRBuilder2 contained 704 experimentally resolved TCR structures sourced from STCRDab [29, 35]. These structures were obtained from 463 distinct PDB files, and 349 of the 704 structures are unique in sequence.

We extended this training set with 204 Immunocore TCR structures, of which 201 sequences are unique and only 6 overlap with the existing training set. Further filtering for alpha/beta TCRs yielded a total of 881 structures, of which 524 are unique in sequence.

TCRBuilder2 was evaluated on a relatively small test set of 21 structures [51]. Since the original work was published more TCR structures have become available, allowing for the curation of a larger and more robust test set. We updated the database of TCR structures, STCRDab [29], and generated a new, independent test set with the following criteria: TCRs must be paired alpha-beta chains, have lower than 3.5Å resolution, be unique within the test set and be non-identical to a paired sequence in the training set. This process yielded 45 TCR structures, the PDB codes of which we have provided in the Supplementary Information (SI Tab. 1).

### 4.2 TCRBuilder2+ model and training

We retrained the deep learning model TCRBuilder2 [35] on our new dataset to create TCRBuilder2+. Like the original model, the neural network architecture of TCRBuilder2+ is based on the ‘structure module’ of Alphafold [34], however, because TCRBuilder2 followed from ABodyBuilder2, in the original work the model ensemble was selected to minimise the RMSD over the beta chain, whose genetic recombination is analogous to the heavy chain. We adapted the ensemble selection in TCRBuilder2+ to account for the structural diversity of the alpha chain by selecting the ensemble of models that minimises the RMSD over both the alpha and beta chain on the validation set. Specifically, ten models were independently trained, each with a partially randomised validation set. The validation sets each contained 30 TCRs which were held constant across all ten splits as well as 20 randomly selected TCRs. After training all ten models to convergence, we evaluated them on the constant subset of the validation set to select the four models with the lowest total RMSD over both the alpha and the beta chain. During inference we predicted four structures and selected the model closest to the ensembled average. The selected structure was then refined using the OpenMM implementation of the AMBER14 protein force field and returned as the final prediction [76, 77]. To produce an uncertainty estimate, we also calculated the sum of the squared distance of each model’s prediction *x_i_* to the mean prediction 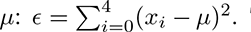. This provides an uncertainty proxy for the coordinates of every amino acid. To produce multiple conformations for both the molecular dynamics and docking simulations we returned all four models’ predictions, and minimised each of them with the AMBER14 force field.

### 4.3 TCR predictions with Alphafold Multimer

We used an instance of Alphafold Multimer [56] installed on our servers, and used version 2.3.1 of the weights. Due to constraints on the memory of the installation, we used the reduced version of the BFD database, as provided accompanying the Alphafold weights. We set a cut-off date of 2020/12/30 for templates, as this excludes all the test structures. However, it is possible that some of the test structures were included in the training data for Alphafold Multimer version 2.3.1, potentially giving the model a slight advantage.

### 4.4 Sequence numbering and gene annotation

We used IMGT numbering and gene annotations computed on amino acids by ANARCI [78, 79]. Table 2 shows the definitions of loops and anchors used. We refer to the CDR3 loops of the alpha and beta chain as CDRA3 and CDRB3 respectively.

### 4.5 Calculating structural deviation

We used the root-mean-squared-distance (RMSD) as the evaluation metric for the predictions, which was calculated over the amino acid backbone atoms (N, C*_α_*, C, and O). We aligned each CDR by the backbone atoms of the anchor residues and calculated RMSD over the backbone atoms of residues in the loops. The loop and anchor residue numbers are provided in Tab. 2, and were annotated using ANARCI [79] according to the IMGT numbering scheme [78].

To compare CDR3 loop structures of different lengths, we implemented spline up-sampling, the parameters of which we set such that the distance calculated is RMSD equivalent when the loops are of the same length. Further details can be found in the Supplementary Information (SI Sec. 5).

An RMSD threshold for an acceptable TCR structure prediction was based on our observations of the sequence to structure identity distributions of CDRA3 and CDRB3 (Fig. 1d & e). Since RMSD will naturally be greater for longer CDR3 loops due to the increased degrees of freedom in the backbone, we also investigated the background distribution within each loop length (SI Fig. 5).

This confirmed 2.0Å as a sensible threshold across the range of TCR CDR3 lengths.

### 4.6 Sequence identity

We calculated the pairwise sequence identity between sequences with the following formula:

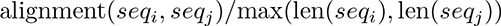

We used alignment(.) as implemented in *Biopython* as *pairwise2.align.globalxx* [80]. The function returns the sum of the alignment of two sequences where the alignment contains 1 for every matched pair of amino acids and zero for every mismatched amino acid pair. No gap or insertion penalties were used.

### 4.7 Clustering

To cluster CDR loops by structure and sequence we implemented a greedy clustering algorithm which we applied to pairwise distance matrices (Alg. 1). For structures the distance matrices contained RMSD between structures, and upsampled RMSD in cases where loop lengths were different. For sequences we calculated a sequence distance matrix from the sequence identity matrix as *d_i,j_* = 1 *− i_i,j_*.

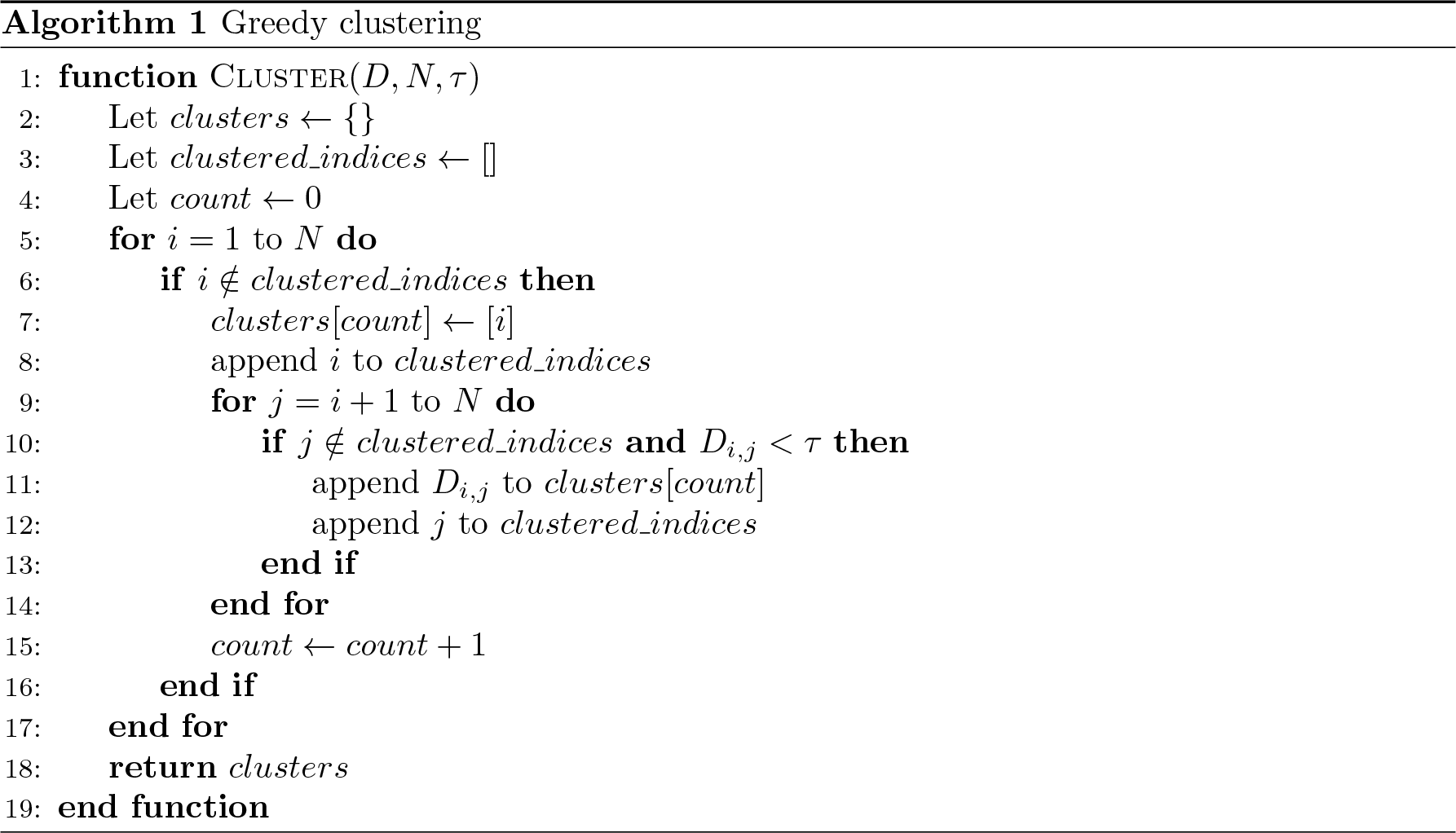

To visualise clusters we used the t-SNE dimension reduction algorithm [81] as implemented in *scikit-learn* [82], with a perplexity parameter of 10. We used the t-SNE plots for visualisation purposes only and used the clustering algorithm as outlined above to colour the clusters.

### Data Availability

TCRBuilder2+ weights: https://doi.org/10.5281/zenodo.10892159 OTS TCR structure predictions made with TCRBuilder2+: https://doi.org/10.5281/zenodo. 10854757.

### Code Availability

TCRBuilder2+ github (integrated into ImmuneBuilder): https://github.com/oxpig/ImmuneBuilder

## Author contributions

N.P.Q, M.I.J.R., S.H., and C.M.D conceived the project and designed the study with input from all authors. N.P.Q. trained the final TCRBuilder2+ model, optimised the model version, conducted the analysis, and compiled the results. B.A. designed and implemented the original ImmuneBuilder model and provided training code for TCRBuilder2+. N.P.Q. compared TCRBuilder2+ against other methods. B.G. conducted and analysed the molecular dynamics study. N.P.Q. ran the docking simulations. V.K. and S.H. provided the additional TCR structure data from Immunocore and advised the project. N.P.Q, M.I.J.R., and C.M.D wrote the manuscript with input from all authors. M.I.J.R, S.H., and C.M.D. supervised the project.

## Footnotes

1 Dataset of natural TCR structure predictions: https://doi.org/10.5281/zenodo.10854757

