## Supplementary Information for "T-cell receptor structures and predictive models reveal comparable alpha and beta chain structural diversity despite differing genetic complexity"

May 20, 2024

#### Contents

**1 SI Tables** **2**

**2 SI Figures** **5**

**3 Accurate prediction of both CDRA3 and CDRB3 is required for TCR:pMHC interface characterisation using physics-driven docking** **14**

**4 Molecular dynamics suggests ensembles of models generated by TCR structure predictors may represent alternative, physically plausible apo states** **18**

**5 Structural distance calculation for length-mismatched CDR loops** **25**

### 1 SI Tables

| | | | | | | | | | |
|---|---|---|---|---|---|---|---|---|---|
| 4nhu | 5wlg | 6zkx | 7dzm | 7dzn | 7f5k | 7fje | 7l1d | 7n2o | 7n2q |
| 7n2r | 7n2s | 7n5c | 7n5p | 7na5 | 7ndq | 7ndt | 7nme | 7ow5 | 7pb2 |
| 7pbc | 7pbe | 7pdw | 7phr | 7q9b | 7qpj | 7r7z | 7rdv | 7rk7 | 7rm4 |
| 7rrg | 7s8i | 7sg1 | 7su9 | 7t2b | 7t2c | 7z50 | 7zt7 | 8cx4 | 8d5q |
| 8dnt | 8es9 | 8gvb | 8gvi | 8shi | | | | | |

SI Table 1: PDB codes of 45 TCRs in the test set.

| Length | CDRA1 | CDRA2 | CDRA3 | CDRB1 | CDRB2 | CDRB3 |
|---|---|---|---|---|---|---|
| 5 | 15 | 19 | - | 21 | 5 | - |
| 6 | 64 | 33 | - | 7 | 57 | - |
| 7 | 17 | 59 | - | - | 8 | - |
| 8 | - | 22 | 8 | - | - | - |
| 9 | - | - | 37 | - | - | 24 |
| 10 | - | - | 80 | - | - | 54 |
| 11 | - | - | 121 | - | - | 99 |
| 12 | - | - | 67 | - | - | 136 |
| 13 | - | - | 77 | - | - | 66 |
| Total | 96 | 133 | 390 | 28 | 70 | 379 |

SI Table 2: Number of clusters of TCRBuilder2+ training structures by greedy clustering by RMSD with 1Å clustering threshold. '-' indicates less than 10 loops of that length existed in the training set to cluster. There are approximately as many CDRA3 clusters as CDRB3 clusters, suggesting CDRA3 loops are as structurally diverse as CDRB3 loops. There are substantially more CDR3 loop clusters than CDR1 and CDR2 clusters.

| Model | A | fwa | cdra1 | cdra2 | cdra3 | B | fwb | cdrb1 | cdrb2 | cdrb3 |
|---|---|---|---|---|---|---|---|---|---|---|
| TCRBuilder2+ | 43 | 45 | 37 | 43 | 21 | 42 | 45 | 44 | 44 | 34 |
| TCRBuilder2 | 45 | 45 | 36 | 45 | 23 | 43 | 45 | 45 | 45 | 26 |
| Alphafold Multimer | 44 | 45 | 37 | 43 | 27 | 43 | 45 | 44 | 45 | 32 |
| LYRA* | 35 | 38 | 27 | 36 | 8 | 38 | 38 | 36 | 38 | 20 |
| ABodyBuilder2 | 0 | 0 | 0 | 0 | 6 | 0 | 1 | 0 | 0 | 4 |

SI Table 3: Number of test set predictions per model with RMSD $< 2$Å. TCRBuilder2+ outperforms TCRBuilder2 and Alphafold Multimer for CDRB3, but retrieves fewer CDRA3 structures. *LYRA, the homology method, could only be evaluated on 38 of the 45 test structures for which a structure was generated.

| Model | Compute resource | Mean inference time per TCR | Relative speed up |
|---|---|---|---|
| TCRBuilder2+ | CPU i7 core | 24.4 sec | 261 x |
| Alphafold Multimer | GPU Ampere A100 | 110 min 19.6 sec | 1 x |

SI Table 4: Inference time of TCRBuilder2+ compared to Alphafold Multimer. TCRBuilder2+ offers a 261-times speed up on CPU compared to Alphafold Multimer on GPU, enabling inference of TCR structures at repertoire scale.

| | 0.5 | 1.0 | 1.5 | 2.0 | 2.5 | 3.0 | 3.5 | 4.0 | 4.5 |
|---|---|---|---|---|---|---|---|---|---|
| CDRA1 | 501 | 126 | 45 | 23 | 12 | 9 | 4 | 3 | 3 |
| CDRB1 | 215 | 34 | 9 | 6 | 5 | 2 | 1 | 1 | 1 |
| CDRA2 | 604 | 200 | 84 | 42 | 27 | 17 | 9 | 7 | 7 |
| CDRB2 | 415 | 105 | 45 | 23 | 13 | 8 | 5 | 2 | 2 |
| CDRA3 | 994 | 840 | 410 | 148 | 62 | 22 | 13 | 5 | 4 |
| CDRB3 | 1000 | 828 | 417 | 178 | 71 | 32 | 19 | 11 | 5 |

SI Table 5: Number of clusters for each CDR loop identified for a sample of 1000 OTS TRCBuilder2+ structure predictions for varying RMSD thresholds in Ångströms.

| | 0.5 | 1.0 | 1.5 | 2.0 | 2.5 | 3.0 | 3.5 | 4.0 | 4.5 |
|---|---|---|---|---|---|---|---|---|---|
| CDRL1 | 90 | 31 | 19 | 16 | 11 | 7 | 6 | 5 | 5 |
| CDRH1 | 121 | 19 | 8 | 5 | 2 | 2 | 2 | 1 | 1 |
| CDRL2 | 25 | 9 | 7 | 6 | 5 | 4 | 4 | 3 | 3 |
| CDRH2 | 156 | 36 | 18 | 11 | 9 | 5 | 4 | 4 | 3 |
| CDRL3 | 391 | 131 | 56 | 23 | 12 | 7 | 6 | 4 | 4 |
| CDRH3 | 995 | 880 | 659 | 406 | 208 | 105 | 61 | 39 | 25 |

SI Table 6: Number of clusters for each CDR loop identified for a sample of 1000 OAS ABodyBuilder2 structure predictions for varying RMSD thresholds in Ångströms.

| Gene type | TRA | TRB | IGL/K | IGH |
|---|---|---|---|---|
| V | 61 | 77 | 83/109 | 205 |
| D | - | 2 | - | 37 |
| J | 61 | 14 | 11/5 | 9 |
| Potential combinatorial diversity | 3721 | 2156 | 913/545 | 68 265 |

SI Table 7: Number of IMGT [1] catalogued human genes for the TRA, TRB, IGL/K, and IGH loci. TRA and TRB are the TCR alpha and beta loci respectively. IGL/K are the antibody light chain loci, split into lambda and kappa genes. IGH is the antibody heavy chain locus. V, J and D are the variable, joining, and diversity segments respectively. A relatively large number of joining genes have been observed for the TRA locus compared to TRB and IG loci. The large number of TRAJ and low number of TRBD result in comparable potential combinatorial diversity of TCR alpha and beta sequences.

| | CDRA3 | CDRB3 |
|---|---|---|
| TCRBuilder2+ | 3.48 | 3.15 |
| Alphafold Multimer | 2.80 | 2.30 |

SI Table 8: Mean number of conformation clusters per TCR sequence predicted, 1Å RMSD threshold.

| | 1.0 | 0.9 | 0.8 | 0.7 | 0.6 | 0.5 | 0.4 | 0.3 | 0.2 | 0.1 | 0.0 |
|---|---|---|---|---|---|---|---|---|---|---|---|
| CDRA1 | 65 | 65 | 50 | 49 | 30 | 22 | 8 | 7 | 2 | 1 | 1 |
| CDRA2 | 59 | 59 | 50 | 48 | 37 | 20 | 16 | 9 | 6 | 2 | 1 |
| CDRA3 | 1968 | 1705 | 1012 | 450 | 168 | 64 | 32 | 15 | 3 | 2 | 1 |
| CDRB1 | 44 | 44 | 44 | 17 | 16 | 8 | 6 | 6 | 2 | 2 | 1 |
| CDRB2 | 57 | 57 | 47 | 47 | 23 | 22 | 11 | 10 | 3 | 2 | 1 |
| CDRB3 | 1998 | 1938 | 1490 | 758 | 256 | 64 | 22 | 10 | 3 | 2 | 1 |

SI Table 9: Number of sequence clusters emerging from 2000 randomly sampled TCRs from OTS, by TCR loop and sequence identity clustering threshold.

### 2 SI Figures

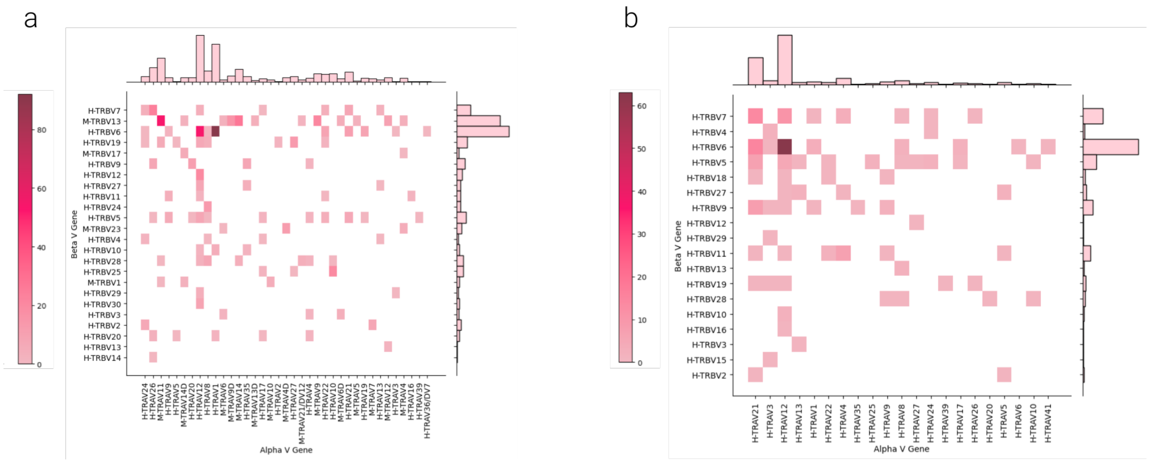


SI Figure 1: Joint distribution of alpha and beta V genes of TCR structures in the training data. a) Joint distribution of TCRs sourced from STCRDab. b) Joint distribution of TCRs provided by Immunocore. Genes beginning with 'H-' are human, those beginning with 'M-' are murine.

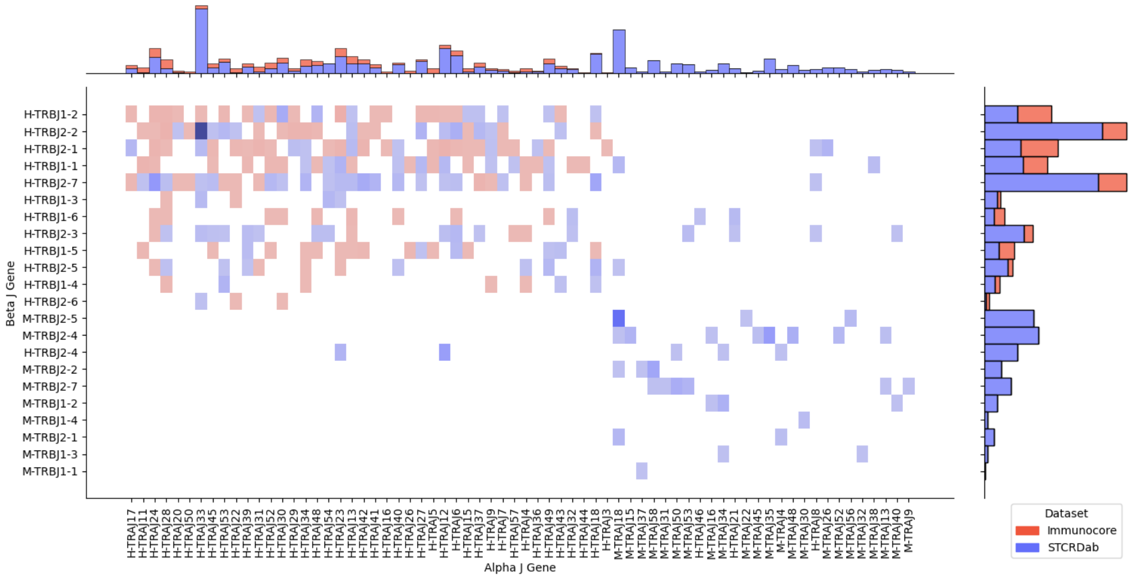


SI Figure 2: Joint distribution of alpha and beta J genes of TCR structures in the training data, coloured by data source (STCRDab in blue, Immunocore in red). Genes beginning with 'H-' are human, those beginning with 'M-' are murine.

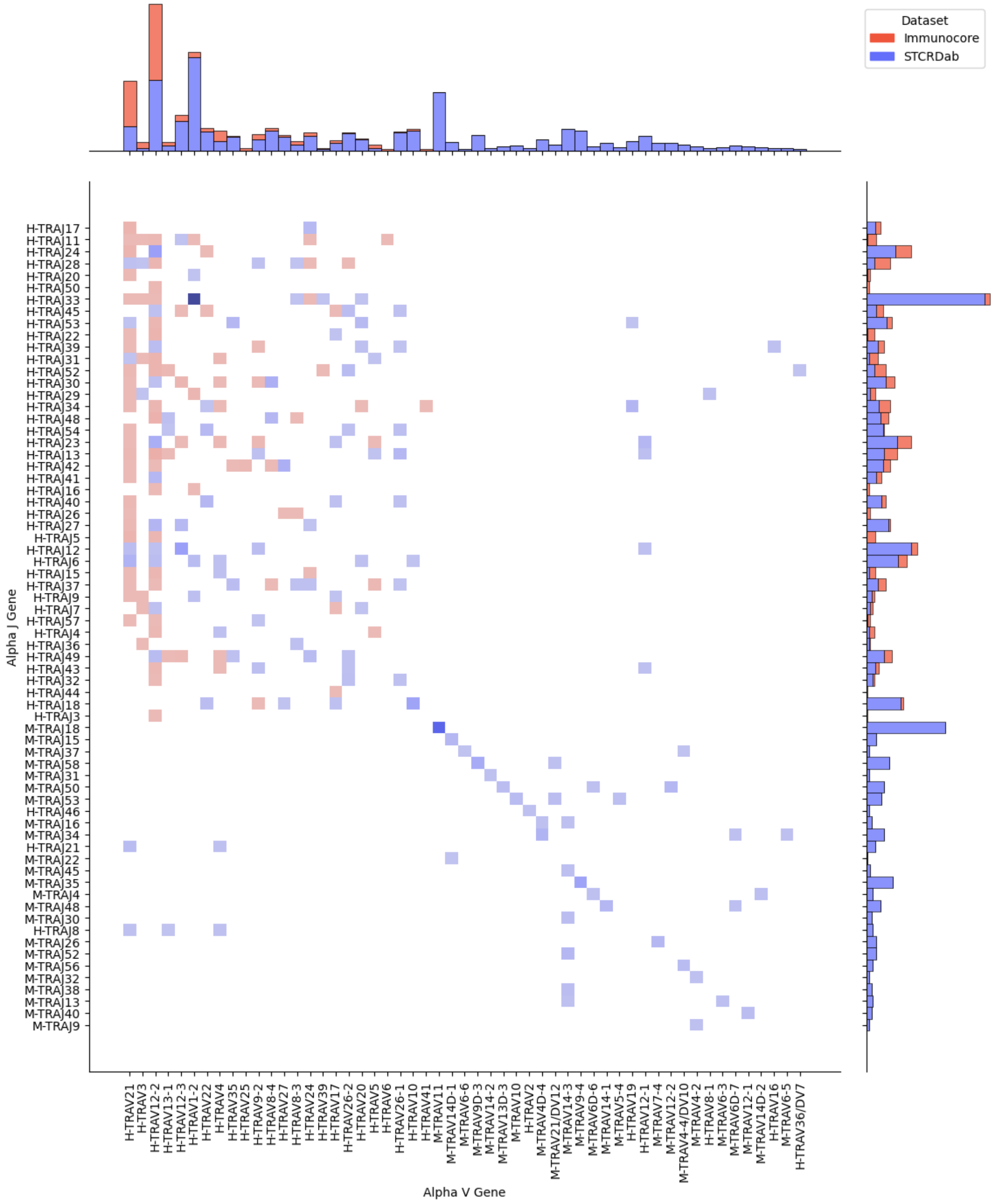


SI Figure 3: Joint distribution of TRAV and TRAJ genes of TCR structures in the training data, coloured by data source (STCRDab in blue, Immunocore in red). Genes beginning with 'H-' are human, those beginning with 'M-' are murine. The TRAV-TRAJ combinatorial space is visibly vaster and more sparsely sampled than the TRBV-TRBJ combinatorial space (SI Fig.4)

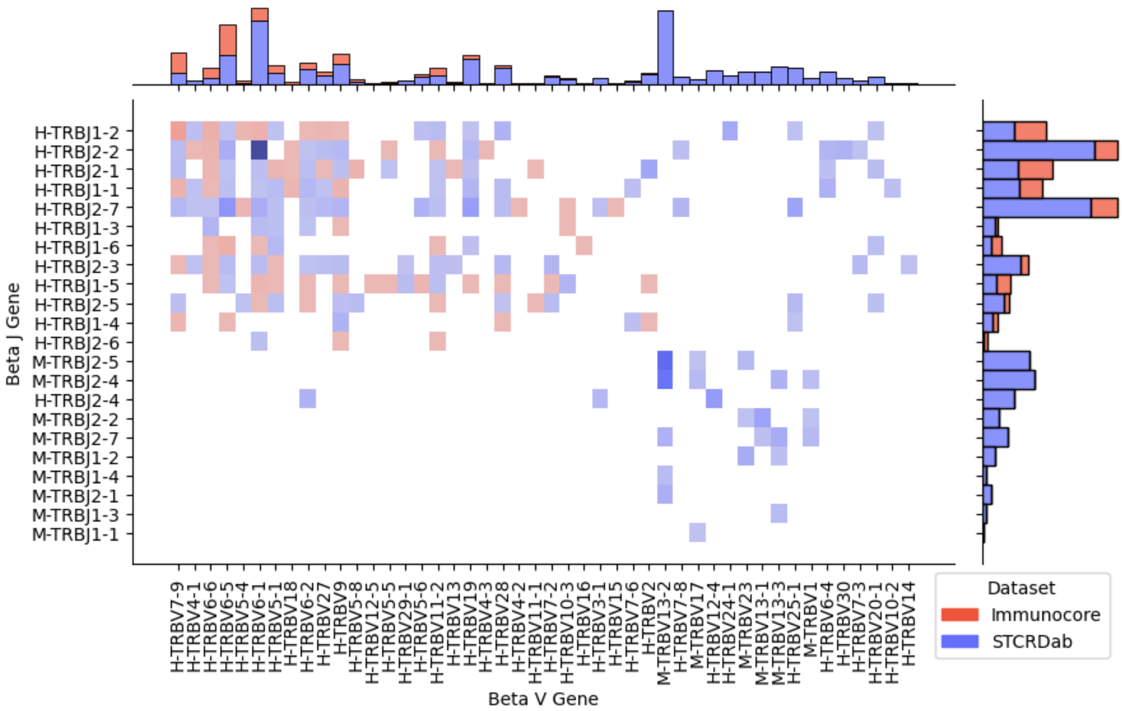


SI Figure 4: Joint distribution of TRBV and TRBJ genes of TCR structures in the training data, coloured by data source (STCRDab in blue, Immunocore in red). Genes beginning with 'H-' are human, those beginning with 'M-' are murine.

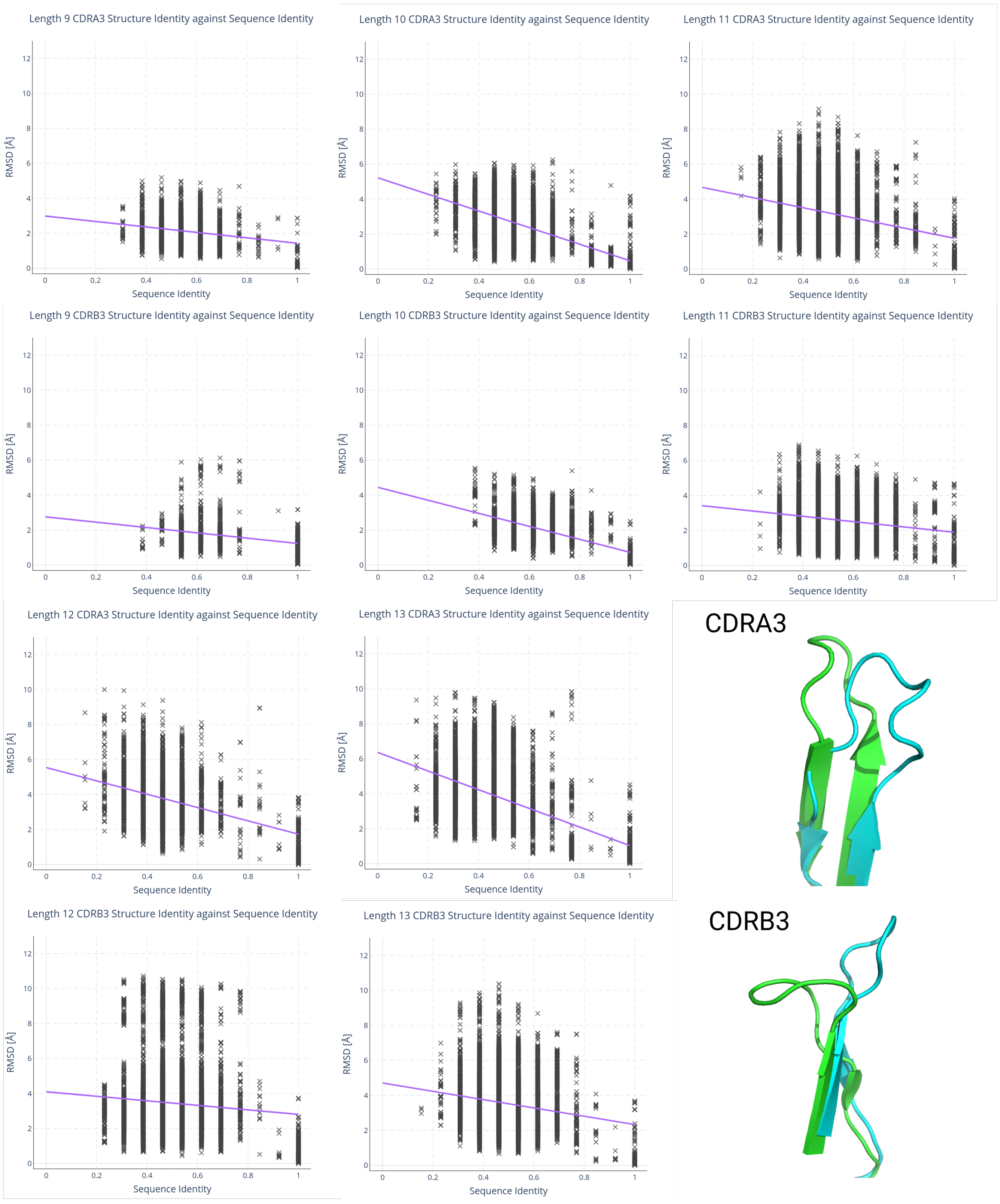


SI Figure 5: Pairwise RMSD *vs* sequence identity distribution of training data by loop length. Examples of identical sequence loops with high RMSD suggest that both CDRA3 and CDRB3 structures are diverse and are often not predictable from just the sequence of the CDR3 region.

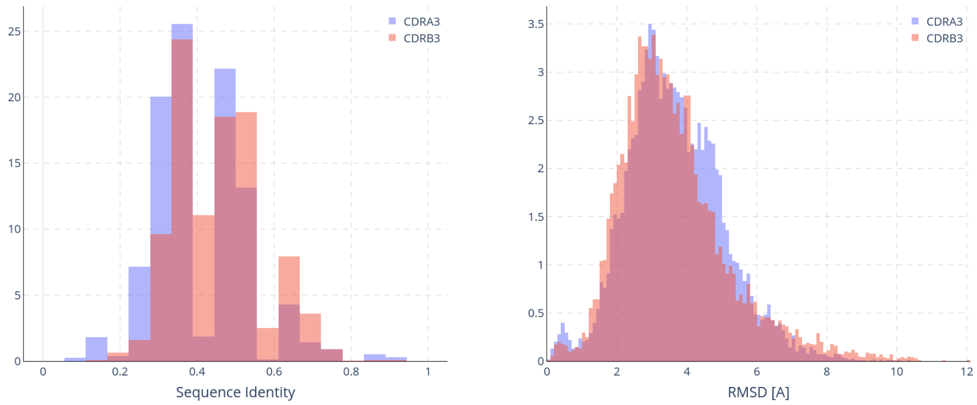


SI Figure 6: Pairwise sequence identity and RMSD (spline upsampled) distributions of the training data of TCRBuilder2+. The distributions of CDRA3 and CDRB3 suggest that CDRA3 is as structurally/sequence diverse as CDRB3.

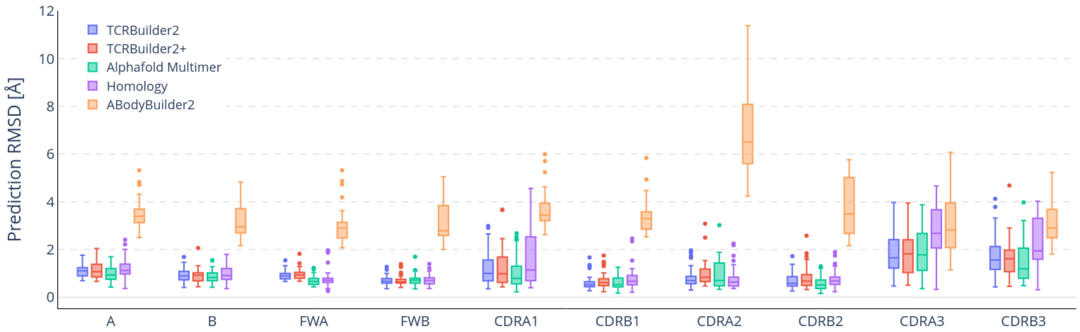


SI Figure 7: Comparison of prediction RMSD by different models, including only the 38/45 test structures for which the homology model LYRA [2] predicted a structure.

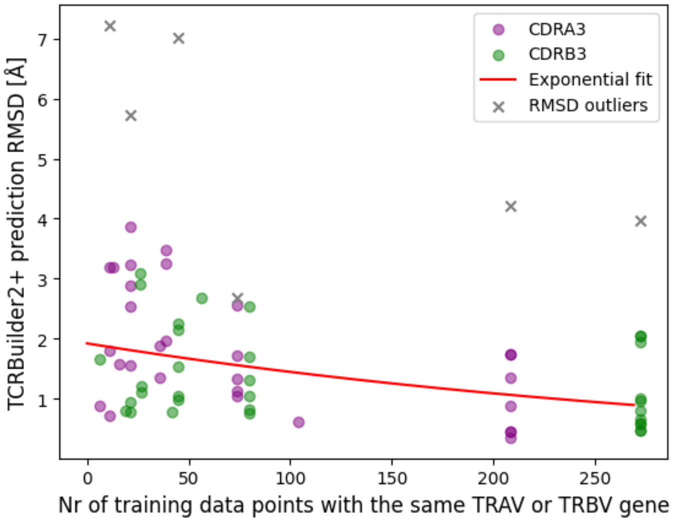


SI Figure 8: Test set TCRBuilder2+ prediction RMSD *vs.* variable gene abundance in training data for CDRA3 (purple), and CDRB3 (green). RMSD outliers of the test structures 7l1d, 7rk7 and 6zkx are shown in in grey. TCRBuilder2+ predictions improve for TCR sequences deriving from a variable gene observed more frequently in the training data. The data support a saturation effect; fitting an exponential curve suggests a slight decay, through the variation about the trend remains high.

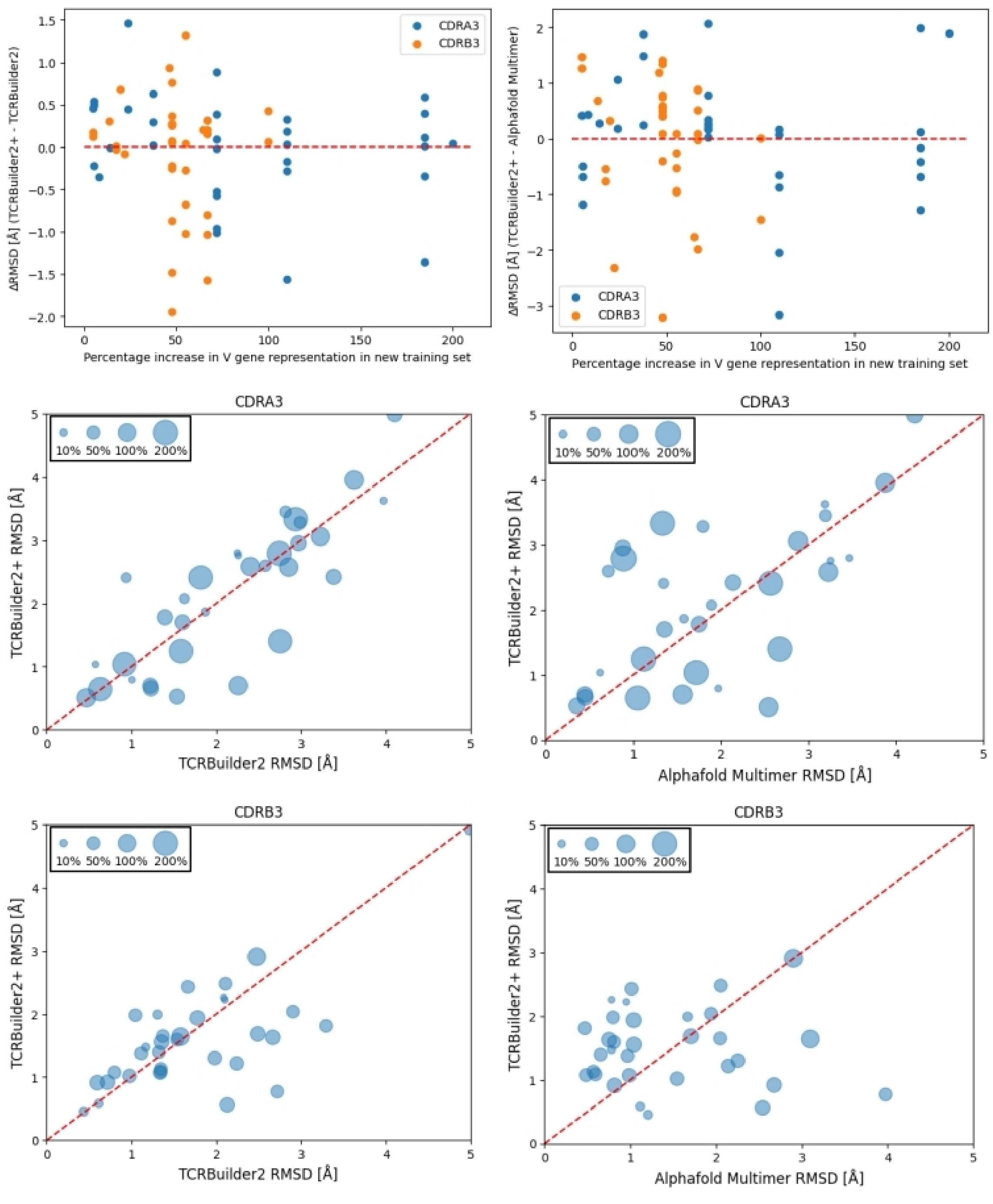


SI Figure 9: RMSD vs relative increase in V gene in training data. a) Difference in RMSD between TCRBuilder2 and TCRBuilder2+ (left) and Alphafold Multimer and TCRBuilder2+ (right) vs percentage increase in V gene in new training set (< 0Å means TCRBuilder2+ is better). b & c) CDRA3 (b) and CDRB3 (c) RMSD of TCRBuilder2+ vs TCRBuilder2 (left) and Alphafold Multimer (right). Bubble size indicates the percentage increase in datapoints for the TRBV gene of a given test sequence in the new training set compared to the old training set, red dashed line is the identity line at which models perform equivalently.

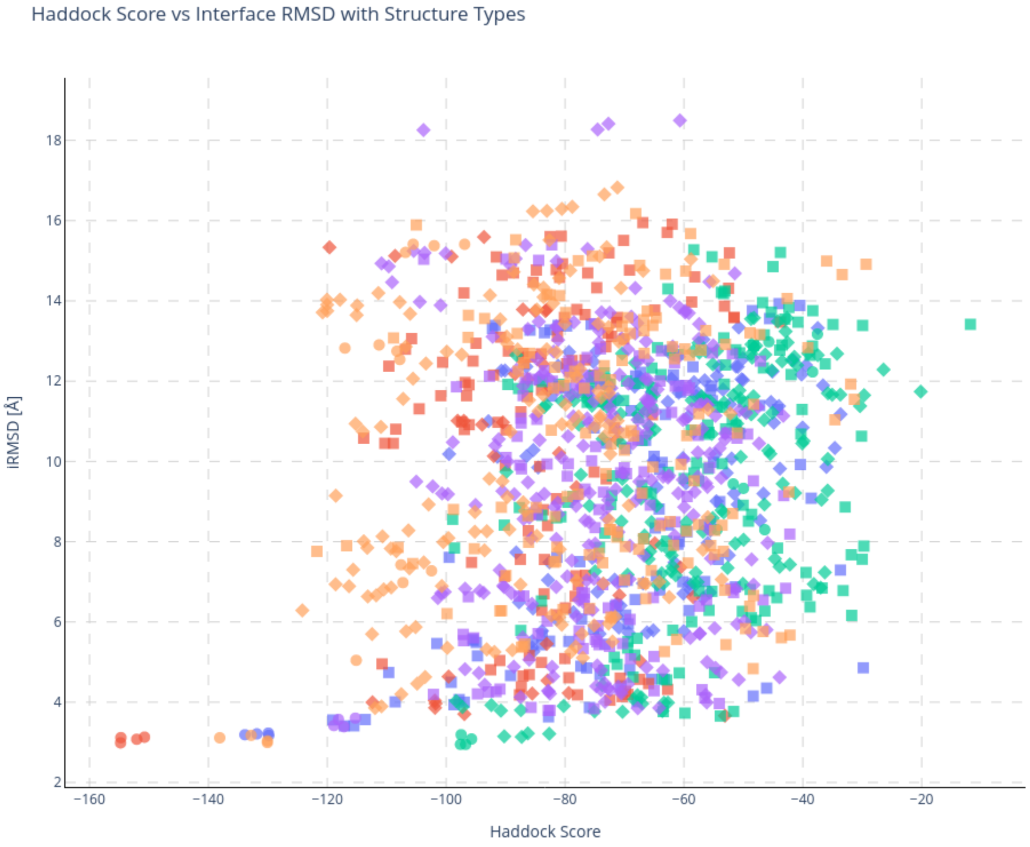


SI Figure 10: All docking clusters iRMSD *vs.* HADDOCK docking score, coloured by PDB code (6zkx: red, 7nme: blue, 7pb2: green, 7rm4: purple, 8d5q: orange). Shapes indicate the type of TCR structure (circle: crystal structure, square: TCRBuilder2+, diamond: Alphafold-Multimer).

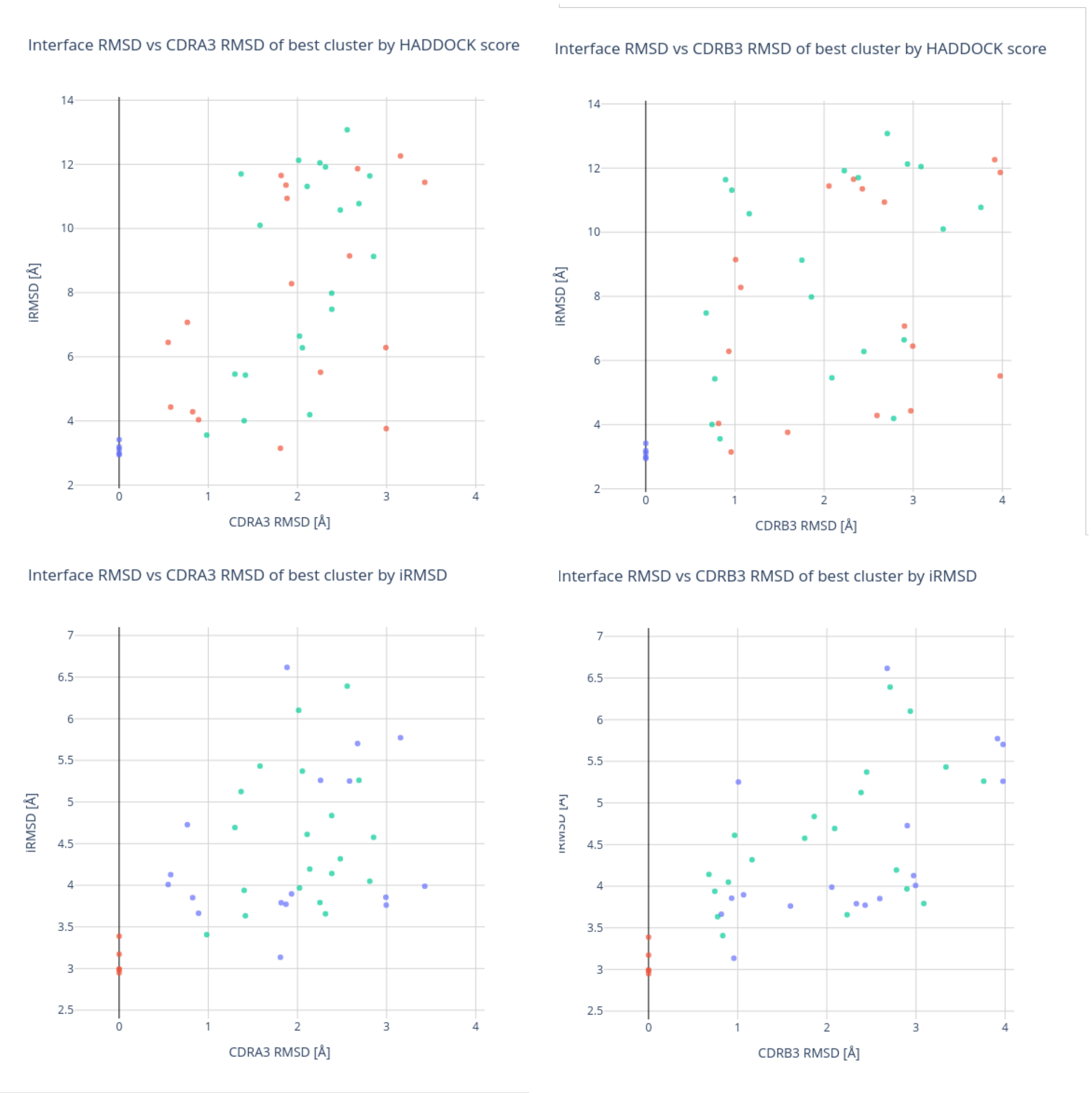


SI Figure 11: iRMSD of best clusters by iRMSD to crystal or HADDOCK score *vs.* CDR3 loop RMSD of TCRBuilder2+ and Alphafold Multimer predictions.

### 3 Accurate prediction of both CDRA3 and CDRB3 is required for TCR:pMHC interface characterisation using physics-driven docking

As presented in the main manuscript (Sec. 2.2), we sought to assess the extent to which accurate CDRA3 or CDRB3 loop structure predictions are key to obtaining useful putative binding poses. We used the software HADDOCK [3] to dock a subset of TCR structures from the test set, both crystal coordinates and models predicted by TCRBuilder2+ and Alphafold-Multimer, to crystal structures of their cognate antigens.

SI Fig. 12a shows the lowest interface RMSD (iRMSD) achieved across the docking run for each model alongside its CDR3 accuracy, while SI Fig. 12b compares CDR3 accuracy to the iRMSD of the 'best' dock defined by docking score. A visual inspection of the docks suggests that an iRMSD of less than 4 Å recaptures the pose with which the CDR3 loops bind to the antigen, with moderate recapitulation of the side-chain interactions (SI Fig. 12c).

The baseline re-docks of the crystal structures of TCRs suggest that the best attainable iRMSD is between 3 Å and 3.5 Å, due to the limited pose sampling of high-throughput docking (SI Fig. 12a & b). Meanwhile, the data on TCR models suggest clearly that accurate prediction of both the CDRA3 and the CDRB3 loops is required to enable accurate docking, defined as iRMSD $<$ 4 Å, particularly when relying on docking score to retrieve the best poses. Specifically, we find that a total RMSD over the CDRA3 and CDRB3 $<$ 3Å consistently enables the retrieval of a dock with an iRMSD $<$ 4Å. It is not possible to identify a consistent accuracy threshold when considering the CDRA3 or CDRB3 individually (SI Fig. 11). This emphasises that both the CDRA3 and CDRB3 tend to contribute to antigen binding and aligns with our previous finding that CDRA3 loops display similar structure diversity to CDRB3 loops.

Though we were able to retrieve TCR:pMHC poses with iRMSD $<$ 4 Å for nearly half of the TCRs, frequently these were not the docks with the best docking score. Quantitatively, while the docking scores of the best ranked dock correlate with iRMSD, only 31.1% (14/45) of the best docks by iRMSD also had the optimal docking score. Furthermore, as the docking scores are calculated as weighted sums of energy terms derived from the docked structure and vary across simulations it is not possible to determine a threshold for docking scores that holds for all TCRs and pMHCs

(SI Fig. 10). The unreliability of general pose ranking functions in this domain limits docking's practical applicability for investigating TCR:pMHC complementarity.

Finally, we investigated whether using multiple distinct TCR structure predictions for each sequence improves docking outcomes; TCRBuilder2+ generates four predictions per sequence (Fig. 1a) and Alphafold-Multimer generates 25, from which one is selected at run time as the best-ranked model. When we docked these alternate structure predictions, we find that the best-ranked TCRBuilder2+ model achieves the lowest iRMSD dock in three of five case studies, while this is only the case for one of five for Alphafold-Multimer. We hypothesise that this is due to the combinatorially expanded search space when including multiple conformations, thereby increasing the chances of retrieving a good docked pose. In the absence of a crystal structure, incorporating multiple distinct TCR conformation predictions in a pipeline may enable the retrieval of better docks, although we note again that the limitations of docking scores mean that identifying the best pose across runs remains challenging.

#### Molecular Docking Methods

We used HADDOCK [3] to run docking simulations. The active residues of the paratope were defined as residues of the six CDR loops according to IMGT numbering [4] and the active residues of the epitope were defined as the residues of the peptide presented by the pMHC. Unburied residues within 6.5Å of the peptide residues are defined as passive residues of the interaction site. No residues are defined as passive in the paratope.

We selected the TCR crystal structures in 6zkx, 7nme, 7pb2, 7rm4, and 8d5q for the docking simulations, as they are crystallised in complex with pMHC and their TCRBuilder2+ and Alphafold Multimer CDR3 prediction accuracies span a range of RMSD values. For each structure we ran docking simulations of the original crystal TCR structure and every TCR structure prediction by TCRBuilder2+ and Alphafold Multimer that clustered into different CDRA3 or CDRB3 conformations by greedy clustering with a threshold of 1Å. Each simulation results in 200 docked poses ranked with HADDOCK's docking score reflective of the enthalpy of the docked interface, clustered by RMSD to the best ranked dock.

We also evaluated docked pose quality by the interface RMSD (iRMSD) to the original crystal structure. iRMSD is calculated as the RMSD between all heavy atoms in the interface after aligning

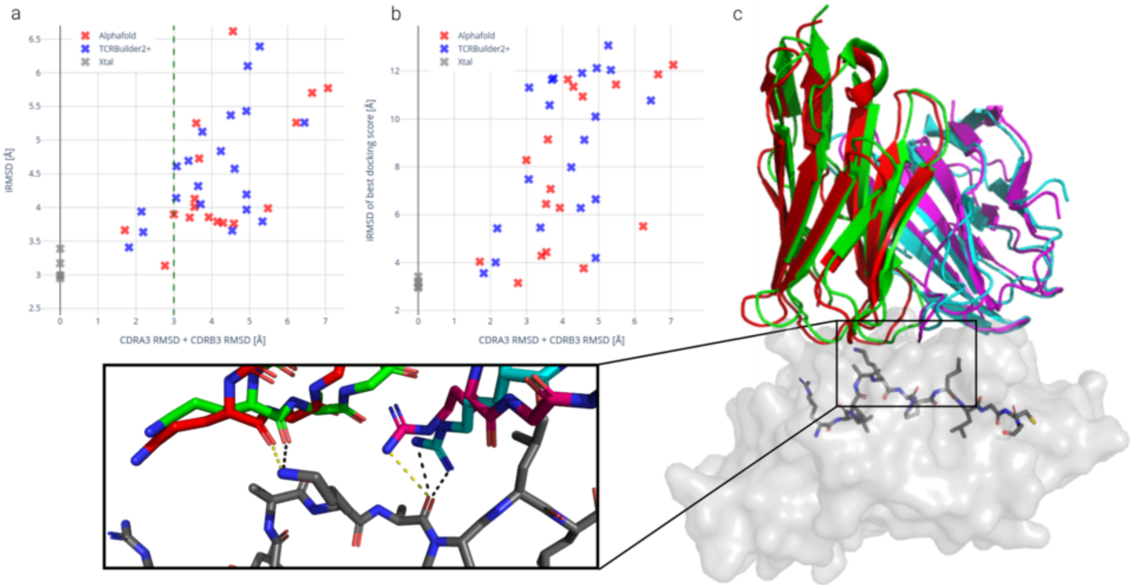


SI Figure 12: a) Best interface RMSD (iRMSD) of docked TCR structures to their cognate antigen. b) iRMSD of the best ranked dock by docking score. Crystal structure re-docks (grey) provide a baseline of realistically attainable iRMSD. Docking iRMSD clearly correlates with CDR3 loop prediction RMSD, with a joint RMSD <3 Å consistently retrieving a dock with iRMSD <4Å. Docking scores do not consistently retrieve the best docked score. c) Docked TCRBuilder2+ structure prediction of 6zkx to crystal structure pMHC. Predicted $\alpha$ and $\beta$ chains are red and magenta respectively, original crystal structure $\alpha$ and $\beta$ chains are green and cyan respectively. The pMHC is shown as a surface with the peptide as sticks. The docked CDRA3 and CDRB3 chains (zoom box) recapture the polar interactions with the peptide (docked interactions in yellow, crystal structure interaction in black).

the dock to the crystal by the MHC backbone atoms (N, $C_\alpha$, C, and O). The interface was defined as all residues of the TCR and pMHC for which the $C_\alpha$ atoms were within 8Å.

### 4 Molecular dynamics suggests ensembles of models generated by TCR structure predictors may represent alternative, physically plausible apo states

TCR CDR3 loops have been hypothesised to be exhibit structural flexibility, particularly when compared to antibody CDR3 loops [5–9]. This provides a challenge to any method that offers a single TCR CDR3 loop conformation prediction and could disproportionately limit expected accuracy. In the main manuscript, we primarily benchmarked the 'top-ranked' predictions of TCRBuilder2+ and Alphafold-Multimer. However, these deep learning models can return multiple predicted structures, which we can analyse to assess whether they could represent physically plausible sets of conformations. TCRBuilder2+ is comprised of an ensemble of four models, each trained with different training and validation splits, and therefore produces four predictions for each sequence. Alphafold-Multimer uses an ensemble of five models which are fine-tuned in two stages from the best performing pre-trained model with different random seeds [10], each of which makes five different predictions for a single TCR sequence, resulting in 25 structures.

We began by investigating the conformational space sampled by the TCRBuilder2+ and Alphafold-Multimer predictions and find that TCRBuilder2+ explores more distinct conformations, despite Alphafold-Multimer producing over six-fold more models. We used greedy clustering with a threshold of 1Å RMSD to investigate the array of conformations predicted by TCRBuilder2+ and Alphafold Multimer (SI Fig. 14) and found that TCRBuilder2+ usually predicts four distinct conformations for both the CDRA3 and CDRB3 loop, while Alphafold Multimer predictions typically fall into just one or two clusters.

In line with our finding that CDRA3 loops are at least as structurally diverse as CDRB3 loops, both models predict a greater number of conformational clusters for the CDRA3 than the CDRB3 (SI Fig.14, SI Table 8). It is unfortunately impossible to disentangle whether this is due to greater uncertainty of the model predictions because the CDRA3 is inherently more difficult to predict, hence leading to less conserved predictions by both models, or whether this is due to the CDRA3 being intrinsically more flexible leading to the prediction of more diverse conformations. In the absence of experimental data that captures flexibility in TCR structures, we use molecular dynamics to assess the physical plausibility of the multiple predicted conformations, thereby evaluating whether predicted structures can be categorised as incorrect. Analysis of these simulations is consistent with more than one CDR3 loop conformation being physically plausible, thereby both supporting the hypothesis that CDR3 loops of TCRs may be flexible [6, 7, 9, 11], and questioning whether regressing structure prediction models towards single crystal structures leads to useful *in-vivo* structure predictions.

Due to the computational expense of running molecular dynamics simulations, we examined the case study of the *apo* crystal structure 7f5k, which we select on the basis that it is not in complex with an antigen, is contained in our filtered test set (SI Table 1), and exhibits two distinct conformations for both the CDRA3 and CDRB3 in the crystal structure(SI Fig. 14).

We then selected four predictions with distinct conformations made by both TCRBuilder2+ and Alphafold-Multimer. Since Alphafold-Multimer makes 25 predictions, we use the four structures that ranked best by pTM score and fall into different conformation clusters. This results in a final set of 10 structures for the same TCR sequence with which we initialise molecular dynamics simulations (SI Fig. 13).

SI Fig. 15 shows the free energy landscape sampled by the simulation trajectories. After being initialised from their predicted or crystal conformation (SI Fig. 13), the CDR3 loops explore the local conformation space, but no macro transitions away from the initial predictions are observed. This suggests that while the predictions do not recapture the loop structure of the crystallised TCR, the predicted structures are energetically stable within the MD forcefield. This supports the hypothesis that multiple loop conformations may be valid, particularly for apo (unbound) TCRs, and that, despite neither model being trained to predict multiple conformations, both Alphafold-Multimer and TCRBuilder2+ could be capturing multiple plausible structures.

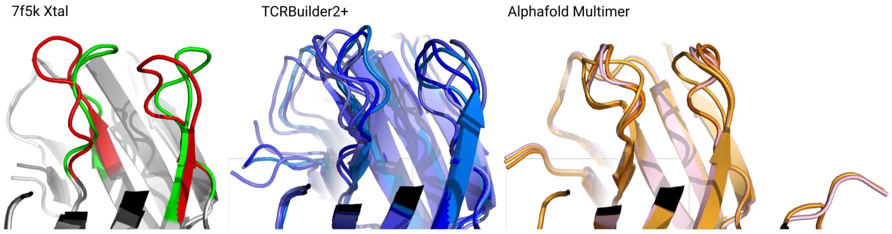


SI Figure 13: Starting conformations used in molecular dynamics (MD) simulations. The 7f5k crystal structure has two distinct conformations (red and green) for both the CDRA3 and CDRB3 loop. TCRBuilder2+ predicts four distinct conformations with which we initialised MD simulations. Alphafold Multimer makes 25 predictions, however many of them are structurally indistinguishable and cluster into a handful of predictions (see SI Fig. 14b). Here, we identified the predictions ranked 0, 1, 14, and 22 as structurally distinct and used them as starting structures for MD simulations.

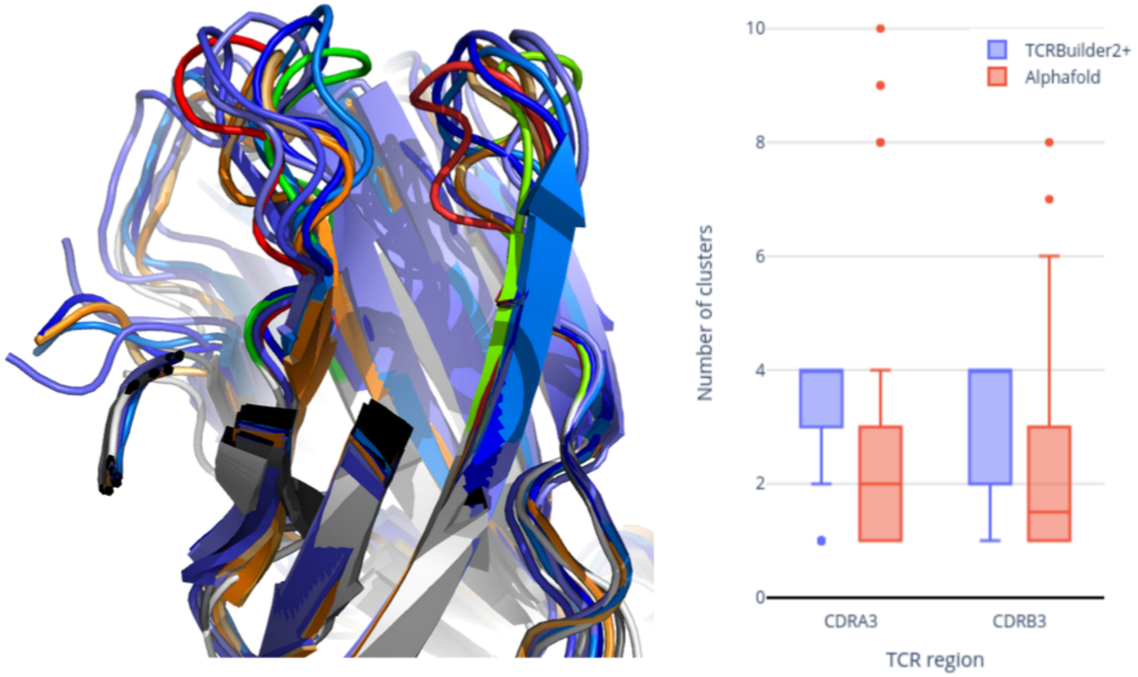


SI Figure 14: a) Multiple CDR3 conformations of 7f5k crystal structure and predicted structures in sliced view. CDRA3 loops are on the left and CDRB3 loops are on the right. The green and red loops are the two conformations observed in the apo TCR crystal structure. The blue loops are the four TCRBuilder2+ predictions, the orange loops are the top four ranked Alphafold-Multimer predictions. The top four Alphafold-Multimer predictions model the same loop, whereas TCRBuilder2+ produces distinct conformations. Neither TCRBuilder2+ nor Alphafold-Multimer predictions recapture either of the crystal structure conformations, but certain trends, for instance the 'inverted S' shape at the base of the CDRA3 loop, are recapitulated. Molecular dynamics simulations suggest that the predicted conformations are physically plausible (SI Fig. 15) and lie within the space of commonly sampled backbone structure for CDR3 loops of this length. b) Number of distinct CDR3 loop structure clusters per sequence predicted for a cluster threshold of 1Å RMSD. Despite TCRBuilder2+ making four predictions, compared to Alphafold-Multimer's default of 25 predictions, the TCRBuilder2+ predictions typically fall into more clusters. Both models predict more conformations for CDRA3 than CDRB3 on average.

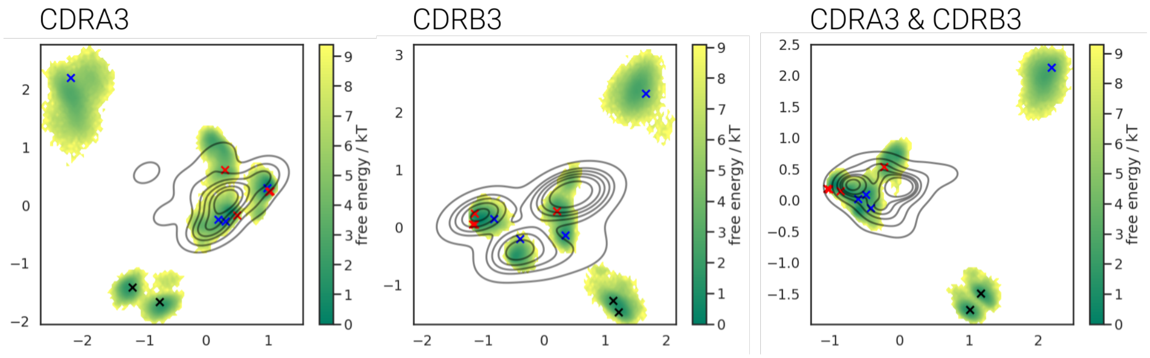


SI Figure 15: Time-lagged independent component analysis (tICA) projections in two dimensions of molecular dynamics trajectories of crystal and predicted CDR3 structures of 7f5k. The structures from which the simulations were initialised are marked as 'x'. The two crystal structure conformations are in black, the four TCRBuilder2+ predictions in blue, and the Alphafold-Multimer predictions in red. The green shading is the free energy distribution calculated from the states sampled in the MD trajectories. The black topography lines are the projected distribution of CDR3 structures of the same loop length as 7f5k. Predicted loop structures occupy spaces with nearby local free energy minima, indicating that the predicted conformations are relatively stable. TCRBuilder2+ samples a greater range of confomational space than Alphafold-Multimer, the crystal structure CDR3 loop structures lie beyond the modes of the background distribution.

#### Molecular dynamics methods

We performed unbiased molecular dynamics simulations on the TCR in PDB entry 7f5k, as it displays multiple distinct conformations for both the CDRA3 and CDRB3 in the crystal structure and was not included in the training set. MD was run both on the crystal poses and on models of the TCR sequence generated by TCRBuilder2+ and AlphaFold-Multimer,

All systems were prepared, and simulations performed, using OpenMM v7.7 [12]. N-methyl groups were used to cap C-termini using an in-house script. Next, using pdbfixer [12], we protonated the models at a pH of 7.5, soaked them in truncated octahedral water boxes with a padding distance of 1 nm, and added sodium or chloride counter-ions to neutralise charges and then NaCl to an ionic strength of 150 nM. Prior to placing molecules in boxes, we aligned all variable region structures on the first copy of the variable region in the 7f5k crystal structure. We parameterised the systems using the Amber14-SB forcefield [13] and modelled water molecules using the TIP3P-FB model [14]. Non-bonded interactions were calculated using the particle mesh Ewald method [15] using a cut-off of distance of 0.9 nm, with an error tolerance of $5 \times 10^{-4}$. Water molecules and heavy atom-hydrogen bonds were rigidified using the SETTLE [16] and SHAKE [17] algorithms, respectively. We used hydrogen mass repartitioning [18] to allow for 4 fs time steps. Simulations were run using the mixed-precision CUDA platform in OpenMM using the Middle Langevin Integrator with a friction coefficient of 1 $ps^{-1}$ and the Monte-Carlo Barostat set to 1 atm. We equilibrated systems using a multi-step protocol detailed in SI Table 10. Following equilibration, we performed 2 µs of unrestrained simulation of the NPT ensemble at 300 K.

We performed dimensionality reduction on the resultant trajectories through Time-lagged independent component analysis (TICA) [19] using PyEMMA2 [20]. We selected the sine and cosine transforms of either CDRA3, CDRB3, or CDRA3 and CDRB3 loop backbone dihedral angles as features, and set 5 ns as the lag time. We also embedded the starting structures for each simulation in the same resultant space. Lastly, to compare other known experimental structures to the conformations observed during simulations, we identified structures of TCR receptors in our training set where both the CDRA3 and CDRB3 loops had the same number of residues as the simulated system (11 residues according to the IMGT definition [4] for either loop) and embedded their CDR3 backbone dihedrals in the same coordinate space.

#### Molecular dynamics equilibration protocol

| Stage | Ensemble | Restrained Atoms | Restraint Strength ($kJ\ mol^{-1}\ nm^{-2}$) | Duration | Start T (K) | End T (K) |
|---|---|---|---|---|---|---|
| 1 | Minimisation | Protein Heavy | 4184.00 | 5000 steps | - | - |
| 2 | NVT | Protein Heavy | 4184.00 | 0.2 ns | 100 | 300 |
| 3 | NPT | Protein Heavy | 4184.00 | 0.2 ns | 300 | 300 |
| 4 | NPT | Protein Heavy | 2092.00 | 0.5 ns | 300 | 300 |
| 5 | Minimisation | Backbone Heavy | 2092.00 | 5000 steps | - | - |
| 6 | NPT | Backbone Heavy | 2092.00 | 0.2 ns | 300 | 300 |
| 7 | NPT | Backbone Heavy | 418.40 | 0.2 ns | 300 | 300 |
| 8 | NPT | Backbone Heavy | 41.84 | 0.2 ns | 300 | 300 |
| 9 | NPT | - | - | 1 ns | 300 | 300 |

SI Table 10: Parameters and steps used during the equilibration of simulated systems.

### 5 Structural distance calculation for length-mismatched CDR loops

Protein structures are typically compared by calculating the RMSD between atom positions, often restricted to $C_\alpha$. However, this poses a problem when comparing chains of different lengths. In order to compare different length CDR3 loop structures, we implement a method for upsampling of the backbone coordinates to match in length. We use spline upsampling to re-sample the shorter loop $C_\alpha$ coordinates by polynomial interpolation, and compare the new coordinates. For chains of the same length this distance calculation is equivalent to standard RMSD. Empirically we find that this method is upper bound by the RMSD, which we find by calculating the distance between two equivalent length loops after upsampling. SI Fig. 16 is an illustrative example of original backbone coordinates (blue) and the upsampled coordinates (red). We also implemented a version of the dynamic time warping (DTW) algorithm, an alternative method based on optimally matching coordinates in the longer loop to coordinates in the shorter loop. While this method is in use, it is liable to spurious, non-RMSD like results when comparing very different structures, such that sequence aligned coordinates are substantially further then DTW match coordinates, resulting in low distances despite structural dissimilarity. We therefore opt for the former method; both methods are implemented as a small python package with a common API (`https://github.com/npqst/different-length-euclidian-rmsd`).

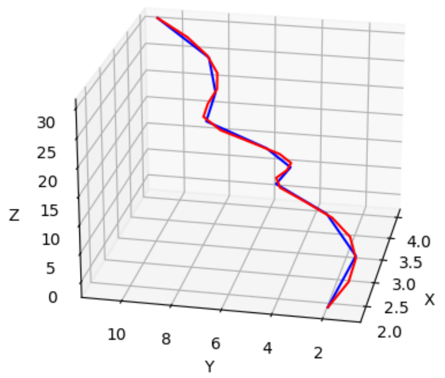


SI Figure 16: Demonstrative visualisation of spline up-sampling on 3D $C_\alpha$ coordinates of a simulated CDR loop. Original coordinates (blue) are up-sampled (red). The coordinates x, y and z are the atom positions in euclidian space.
